# The N-terminal order-disorder transition is a critical determinant for a metamorphosis of IscU

**DOI:** 10.1101/2025.01.09.632086

**Authors:** Jongbum Na, Joongyu Heo, Minchan Jeong, Sangho Ji, Young Ho Ko, Alaleh Shafiei, Nilufer Baldir, Hasan DeMirci, Wookyung Yu, Jin Hae Kim

## Abstract

IscU, a key scaffold protein mediating the biogenesis of iron-sulfur (Fe-S) clusters, exhibits metamorphic characteristics crucial for its versatile and efficient function. Previous studies have demonstrated that IscU has two interconverting conformations: the structured state (S-state) and the disordered state (D-state), each contributing to its distinct functionality and interaction network. Despite its physiological importance, the precise mechanism underpinning the maintenance of IscU’s unique structural heterogeneity has remained elusive. In this study, we used computational and experimental approaches to reveal that the N-terminal order-disorder plays a critical role in the metamorphic modulation of IscU. We found that the N-terminal region displays greater structural plasticity, which is linked to other regions of IscU through coevolutionary relationships. Moreover, we demonstrated that the degree of orderliness in the N-terminal region correlates positively with the stabilization of IscU’s S-state and negatively with its affinity for HscA. This indicates that the flexibility in the N-terminal region is finely tuned to optimize IscU’s physiological efficiency and efficacy. Finally, our data suggest that a peptide mimicking the N-terminal motif can modulate IscU’s metamorphic properties, suggesting a novel therapeutic potential for related pathogenic processes.

## Introduction

The Fe-S cluster is a ubiquitous and indispensable cofactor for various proteins. An imbalance of Fe-S clusters in cells can cause iron accumulation, oxidative stress, and disruption of metal homeostasis, leading to severe human diseases such as neurodegeneration and cancers^1–2^. The Fe-S cluster is synthesized through a few dedicated pathways in the cell. Among them, the Iron-Sulfur Cluster (ISC) system is a primary housekeeping mechanism for maintaining proper levels of Fe-S clusters^3^. The ISC mechanism relies on a tight and complex interaction network with several critical proteins. One key protein is IscU, the scaffold protein responsible for assembling and transferring the Fe-S cluster to other proteins, with assistance from several partner proteins, such as IscS, Fdx, HscA, and HscB^4^.

IscU is classified as a metamorphic protein owing to its unique ability to undergo conformational interconversion between the S-state and D-state under physiological conditions^5^. This conformational heterogeneity can be altered by partner proteins and ligands, which help maximize the functionality of IscU^5^. For example, metal cations, such as Zn^2+^, stabilize the S-state of IscU, which appears suitable to coordinate and sustain the assembled Fe-S cluster. HscB prefers the S-state of IscU to further increase the stability of the Fe-S cluster-bound IscU, while HscA stabilizes the D-state to decrease the affinity for the Fe-S cluster and facilitate its transfer^6^.

Despite the importance of elucidating structural and functional features of IscU, its structural heterogeneity has hampered the consistency of atomic-resolution structural models, particularly at the N-terminal region. Solution nuclear magnetic resonance (NMR) spectroscopic studies have reported that the N-terminal region of IscU is highly disordered^5, 7^. In contrast, the residues S4-N13 in the same region form an α-helix (α1) and directly interact with other regions of IscU (e.g., α-helices at the residues C63-K78 [α2] and the residues K89-E98 [α4]) in the X-ray crystallographic models^8–11^. Notably, previous studies have reported that the N-terminal region of IscU, including α1 and the nearby highly conserved Y3 residue, contributes to constructing the binding interface for the cysteine desulfurase, NFS1 in humans or IscS in *E. coli*^9, 12–13^, suggesting its indispensable roles for IscU’s structure and function. Thus, we speculated that the N-terminal structural flexibility of IscU might be intrinsically maintained and correlated with its metamorphic property.

## Results

### The N-terminal Heterogeneity of IscU Revealed by Computational and Experimental Approaches

To investigate the structural heterogeneity of IscU, we first employed the AF-cluster method with multiple sequence alignment (MSA) of IscU. The AF-cluster method predicts various conformations of a metamorphic protein by running AlphaFold 2 (AF2) predictions based on the clustered MSA data^14^. The superposition of the predicted models showed that the N-terminal region has broader structural divergence compared to other regions of IscU **(Fig. 1a)**. Notably, along with the low prediction confidence at the N-terminal region, the α1 motif was predicted to adopt both disordered and α-helical conformations. These results are consistent with a higher level of structural disorder at the N-terminus of IscU **(Extended data Fig. 1)**^15^.

**Fig. 1.**
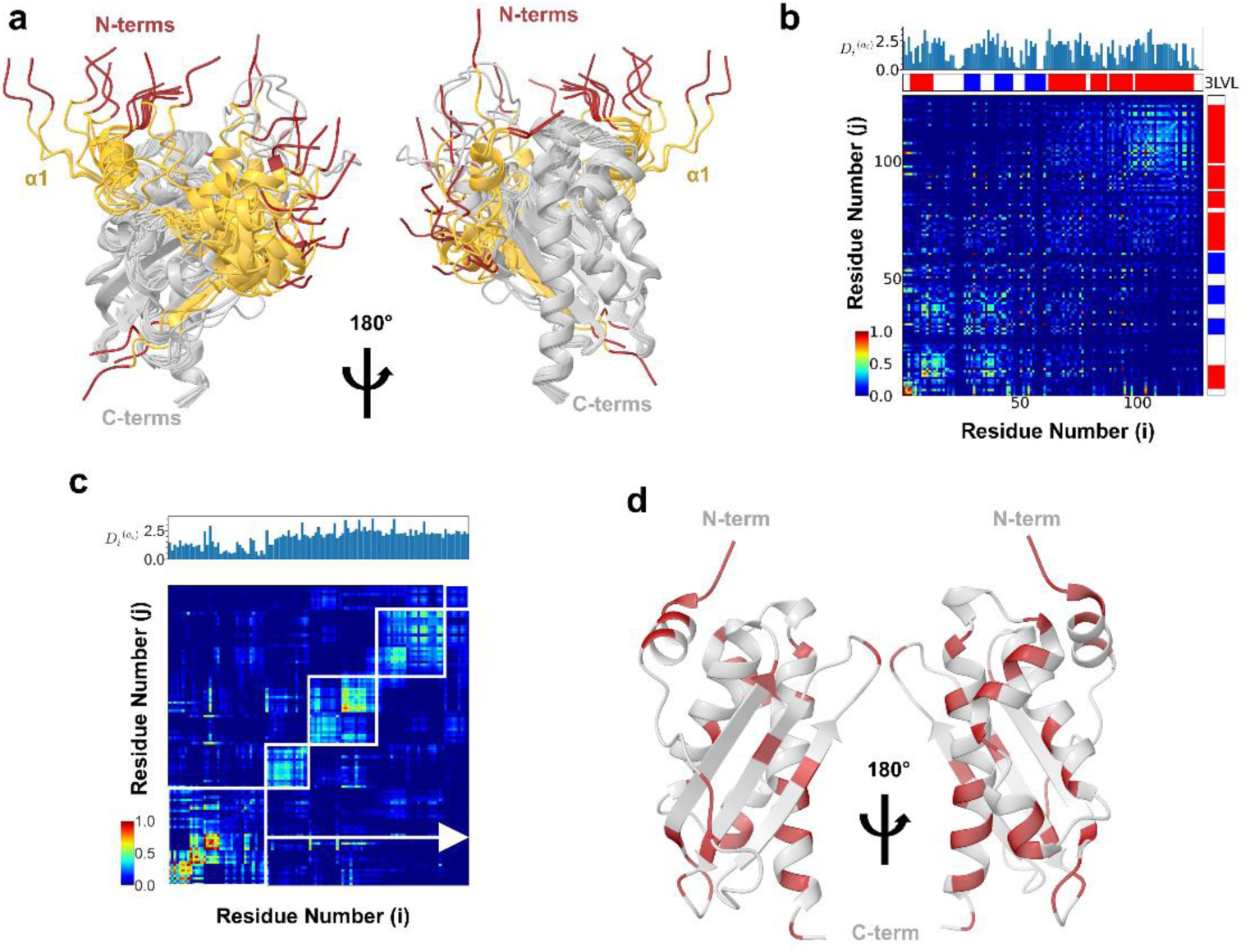
Computational analyses of the heterogeneous N-terminal structures of IscU. **a,** Superposition of AlphaFold 2 predictions of MSA clusters with more than 20 sequences each. Residues at the N-terminus (1-3; N-terms) and the α1-helix motif (4-14; α1) are colored red and yellow, respectively. **b,** Statistical coupling analysis (SCA) of the coevolutionary relationship between amino acid residues of IscU is shown on the representative *E. coli* sequence (UniProt: P0ACD4). Red indicates a high evolutionary relationship between two residues, while blue indicates low coevolution. The secondary structural elements of IscU, which were observed in the IscU-IscS complex model (PDB: 3LVL), are shown for reference. Kullback-Leibler divergence value, 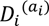, shows the conservation of each residue *i*. **c,** The highly coevolving residues of IscU were clustered using *k*-means and shown as sectors (indicated by boxes) in the plot. **d,** The residues exhibiting a coevolutionary relationship with the residues at the N-terminus and the α1-helix motif are shown in red in the structural model of IscU (PDB 3LVL).

Subsequently, we utilized the statistical coupling analysis (SCA) method to identify a potential correlation between the N-terminal flexibility and metamorphism of IscU. SCA identifies higher-order energetic relationships between residues based on the statistical coevolutionary pattern observed in MSA analysis results^16–18^. Our analysis of the coevolutionary matrix of IscU revealed that residues at the N-terminal tail form a coevolutionary hotspot and coevolve with remote IscU motifs **(Fig. 1b)**. Furthermore, *k*-means clustering results indicated that coevolution partner residues are distributed throughout the entire IscU molecule, implying that the N-terminus of IscU maintains structural heterogeneity. Intriguingly, the Kullback-Leibler divergence (KL divergence) value per residue, 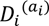, which quantifies the position-specific conservation in the SCA analysis, was lower in the N-terminal region compared to other regions. This suggests the presence of a coevolving but unconserved set of residues in the N-terminal region **(Fig. 1c-d and Extended data Fig. 2)**^16–18^.

To validate the heterogeneous structural states of the N-terminus, we employed a paramagnetic relaxation enhancement (PRE) NMR technique^19^. We created the IscU A2C variant and attached a spin label, a stable nitroxide radical in this study, at the C2 position using the thiol-maleimide reaction **(Extended data Fig. 3 and Supplementary Fig. 1)**^20^. We found that A2C showed mild yet noticeable PRE effects on several regions of proteins **(Fig. 2)**. The residues 16, 18, 20, 24, 31, 33-34, 59, 83, 90, 107, and 112 exhibited more signal decrease than the other regions (**Fig. 2b**). Notably, this result is inconsistent with the structural model with the ordered N-terminus, as it would have resulted in more evident PRE effects on the regions around α3, α4, and α5 **(Fig. 2b)**. Instead, the PRE effects were widespread across several secondary motifs, implying heterogeneous structural states of the N-terminus under our experimental conditions, as analyzed using AF-cluster and SCA (**Fig. 1**). Although AF-cluster cannot give any population-related information for each structural state, we confirmed that the heterogeneous ensemble from AF-cluster was well represented in our PRE data (**Fig. 2**). By comparing the PRE data of IscU A2C with the distance profile (the distances from C_β_ of the N-terminal A2 residue to C_β_ of other residues; **Extended data Fig. 1b**) of AF-cluster ensemble of IscU, we observed consistent patterns in several regions of IscU, i.e., lower PRE values at the regions that were predicted closer to the N-terminus in the AF-cluster ensemble (**Fig. 2a**; highlighted in green and red)^21^. These results indicate that IscU maintains its flexible N-terminus on purpose.

**Fig 2.**
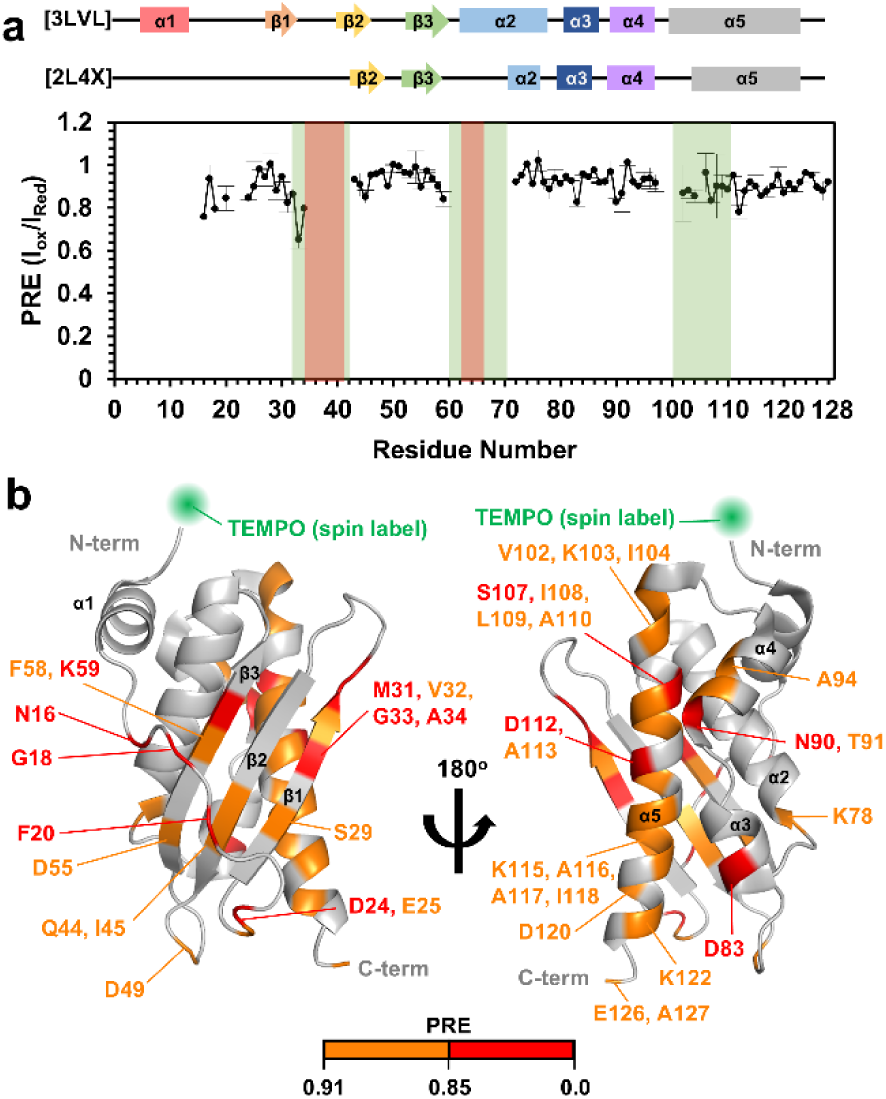
NMR analysis of the heterogeneous N-terminal region of IscU. **a,** Paramagnetic relaxation enhancement (PRE) NMR of IscU A2C with a spin label tag (TEMPO) at the C2 position. PRE(I_ox_/I_red_) of A2C were measured twice, and the averaged values were shown here along with the standard deviations as errors (black). The highlighted regions indicate the residues whose distances from the N-terminal C2 residue in the AF-cluster ensemble were within 25 Å (green) or 23 Å (red) (see Extended data Fig. 1). The secondary structural motifs (from the models with PDB codes 3LVL and 2L4X) are indicated on the top for comparison. **b,** The residues with PREs between 0.91 and 0.85 and the residues with PREs lower than 0.85 are colored orange and red, respectively, on the structural model of IscU (PDB: 3LVL) The mean PRE value across all residues was 0.91 with a standard deviation of 0.06. The green sphere indicates the location of the TEMPO at C2.

### Y3 at the N-terminal Region Is a Critical Residue for IscU’s Structural Heterogeneity

Next, we focused on how structural diversity is maintained at the N-terminal region, starting with Y3, a highly conserved and functionally important residue^9, 12–13^. Some X-ray crystallographic structural models of IscU showed the side chain of Y3 facing the hydrophobic interior of IscU and contacting hydrophobic residues within the α1 motif, suggesting its role in stabilizing the ordered conformation of α1 as well as entire IscU **(Extended data Fig. 4)**^8–10^. To test this hypothesis, we created three IscU mutants, Y3W, Y3M, and Y3A, and investigated their conformational alterations. Size exclusion chromatography (SEC) analyses demonstrated that the effective sizes of Y3 mutants and WT in solution were in the following order: Y3A > WT > Y3W ∼ Y3M **(Fig. 3a and Supplementary Table 1)**. We estimated the helix composition and stability of Y3 mutants using far-UV circular dichroism (CD) spectroscopy, and found that Y3A had fewer α-helical regions with decreased stability against heat, while Y3W and Y3M showed no significant difference compared to WT **(Fig. 3b-c and Supplementary Table 1)**. To investigate the structural perturbation caused by Y3 substitutions at a residue level, we used NMR spectroscopy. For each variant, we collected a series of NMR spectra and assigned the signals corresponding to the S-state of IscU **(Extended data Fig. 5 and Supplementary Fig. 2)**. Chemical shift perturbation (CSP) analyses of Y3 mutants (vs. WT) consistently indicated significant structural perturbations in the β3, α2, α4, and α5 regions of S-state IscU **(Fig. 3d)**. A previous NMR-based study has demonstrated that the intensity ratio between the two signals of K128, corresponding to the S-state and D-state, respectively, in the ^1^H-^15^N heteronuclear single quantum coherence (HSQC) spectrum of IscU, is a reliable indicator to estimate the overall ratio of the S-state and D-state^5^. Upon examining this for Y3 variants, we confirmed that Y3A and Y3W exhibited, respectively, increased and decreased D-state populations **(Fig. 3e and Supplementary Fig. 2d)**. We also analyzed the changes in backbone dynamics caused by Y3 mutations. Compared to WT, Y3A showed a slight decrease in the overall signal intensity in the ^1^H-^15^N heteronuclear Overhauser effect (het-NOE) measurement, indicating increased mobility of the S-state conformation **(Extended data Fig. 6a)**. These results consistently indicate that the residue at the position of Y3 is an important contributor to modulating the S-state and D-state populations of IscU.

**Fig 3.**
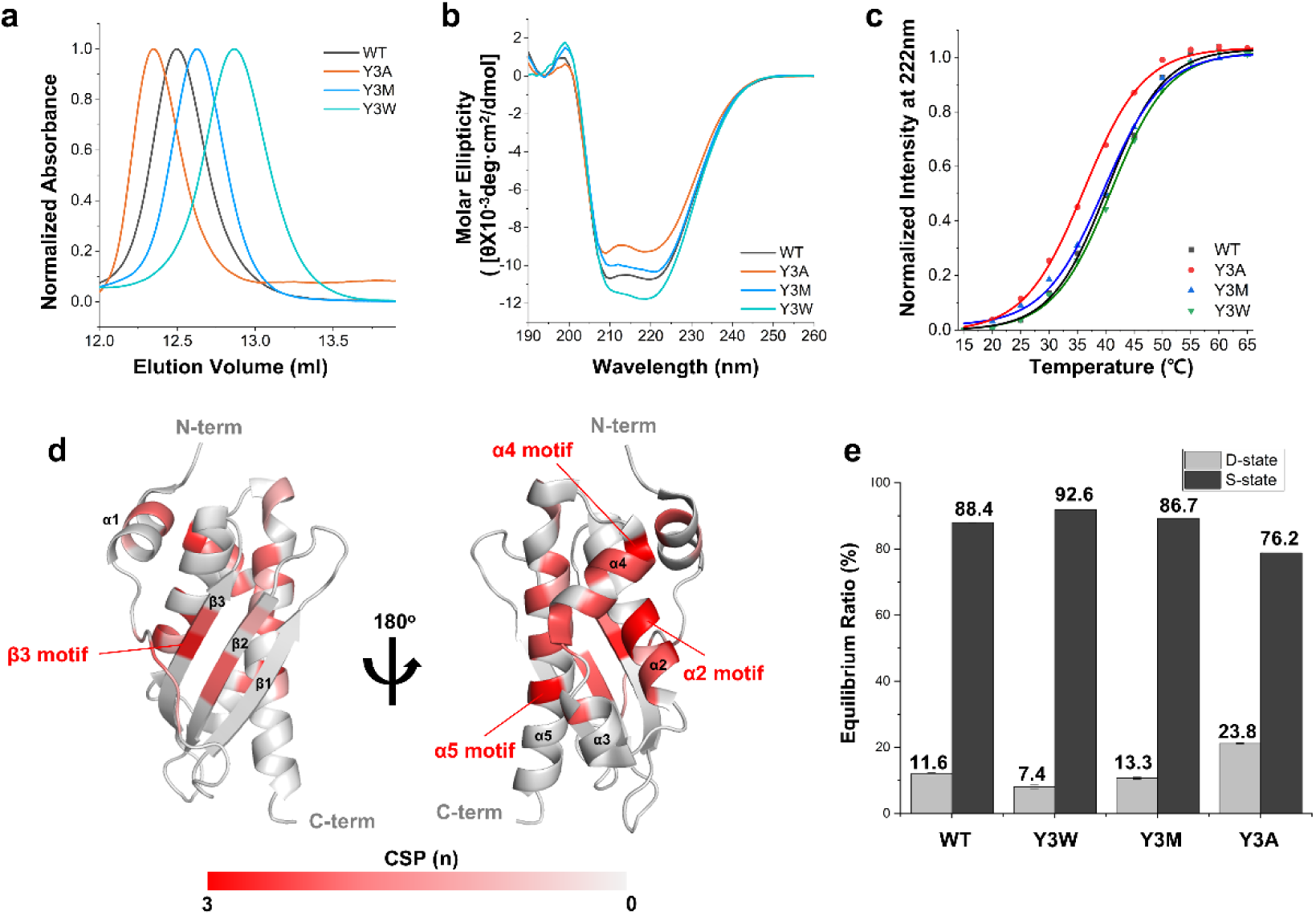
Structural changes of IscU caused by Y3 mutations. Size exclusion chromatography (**a**), far-UV circular dichroism (CD; **b-c**), and NMR spectroscopic data (**d-e**) consistently demonstrated a conformation transition of IscU with Y3 mutation; Y3A induced a less compact, more D-state-like conformation, while Y3W caused a more stable S-state-like conformation. The thermal transitions of IscU variants (**c**) were determined by monitoring CD signals at 222 nm over a temperature range of 15 to 75 °C. The CSP profiles for Y3 variants were calculated by assigning their NMR signals and comparing them to those of WT. Residues showing more frequent perturbations in Y3 variants are shown in more intense red (**d**). A scale of 0 to 3, represented by a color gradient of grey to red, was used, with 3 (intense red) indicating significant perturbation in all Y3 variants and 0 (grey) indicating no perturbation in any Y3 variants (see Extended data Fig. 5 for individual CSP plots and structural mapping results). The ratios of the S-state and D-state (**e**), calculated by comparing the corresponding ^1^H-^15^N NMR signal intensities of the K128 residue, also indicated that Y3W and Y3A stabilize the S-state and D-state, respectively.

### Impact of the N-terminal α-Helix on Structural Heterogeneity of IscU

Subsequently, we questioned whether the N-terminal α-helix (α1) may also affect the structural diversity of IscU. To address this, based on the α1-visible structural models, we designed the IscU variants V7G, D9G, R15A, Y11A, and Y11P, which were expected to hamper the formation of α1 **(Extended data Fig. 7)**. Additionally, we created the IscU variant I8K, where K8 may form a salt bridge with E12 and increase the stability of α1. Our NMR analysis confirmed that I8K had an increased propensity for α1 formation **(Extended data Fig. 8)**. We then employed SEC, far-UV CD, and NMR spectroscopy to investigate the structural changes of these α1 variants. SEC analysis showed that the effective sizes of the variants were in the following order: Y11P > R15A > Y11A > V7G > D9G > WT > I8K, indicating that stable α1 formation increases the S-state population **(Fig. 4a, 4e, and Supplementary Table 2)**. To exclude the possibility that the elution volume change at SEC analysis might originate from a dimer formation, we conducted SEC with multi-angle light scattering analysis and confirmed that even the Y11P variant, which was eluted the earliest at SEC, had a monomeric state under our experimental condition **(Supplementary Fig. 3)**. The CD signal at 222 nm of IscU variants confirmed that the variant with a higher propensity to form α1 stabilized a more α-helical structure **(Fig. 4b and Supplementary Table 2)**. Similarly, based on temperature titration CD results, the transition temperature of I8K was higher than WT, whereas that of R15A was lower, demonstrating a positive correlation between α1 formation and the stability of the S-state conformation **(Fig. 4c and 4e, and Supplementary Table 2)**. The CSP in the ^1^H -^15^N HSQC spectra of the variants compared to WT showed that the β3, α2, α4, and α5 motifs, which were perturbed in Y3 variants, were also significantly affected. We also observed additional CSP in the α1 variants, with the α3 motif experiencing more significant CSP **(Fig. 4d, Supplementary** Fig. 4**, and Extended data Fig. 9)**. The ^1^H -^15^N het-NOE measurements demonstrated that R15A exhibited greater flexibility than WT, whereas I8K had increased stability in its S-state conformation **(Fig. 4e and Extended data Fig. 6)**. All these data demonstrated that the stability of the α1 helix is correlated with the overall conformational diversity of IscU. It also appears likely that the amino acid sequence of α1 is carefully designed and balanced in evolution, which may contribute to keeping the structural and functional flexibility of IscU.

**Fig 4.**
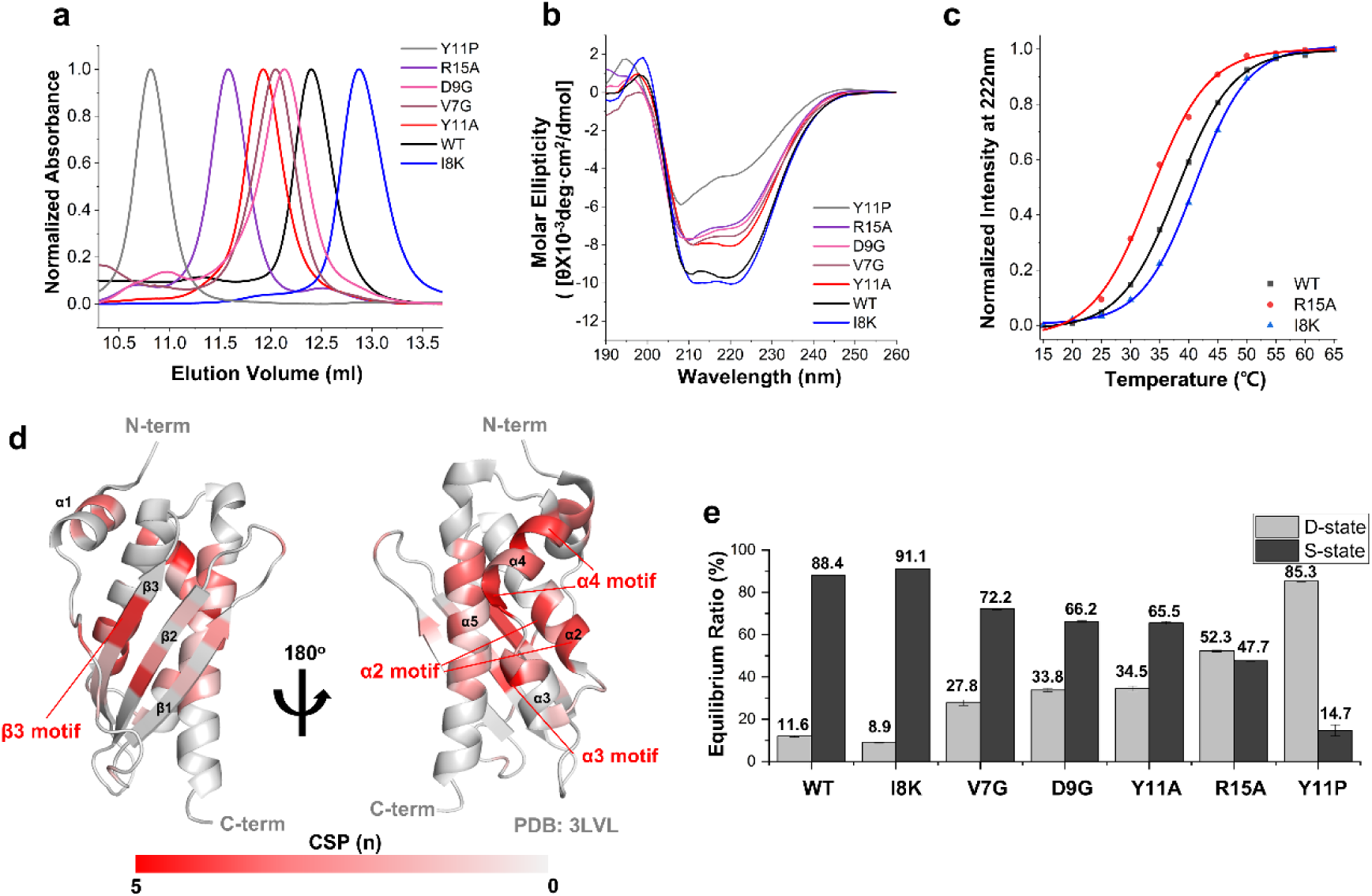
Structural changes of IscU caused by single residue replacements in the α1 helix. Size exclusion chromatography (**a**), far-UV circular dichroism (CD; **b-c**), and NMR spectroscopic data (**d-e**) consistently demonstrated the conformational transition in IscU due to α1-helix mutations. V7G, D9G, Y11A, R15A, and Y11P induced a less compact and unstable conformation, while I8K caused a more stable conformation. The thermal transition of IscU variants (**c**) was determined by monitoring CD signals at 222 nm over a temperature range of 15 to 75 °C. The CSP profiles for α1-helix variants were calculated by assigning their NMR signals and comparing them to those of WT. Residues showing more frequent perturbations are shown in more intense red (**d**; see Extended data Fig. 9 for individual CSP plots and structural mapping results). The ratios of the S-state and D-state (**e**), calculated by comparing the corresponding ^1^H-^15^N NMR signal intensities of the K128 residue, also indicated that I8K stabilizes the S-state, while the others shift the conformational equilibrium towards the D-state. Of note, Y11P exhibited the highest increase in population for the D-state (see Supplementary Fig. 4f).

Based on these results, we reasoned that the α1-like peptide may function as a modulator for IscU’s metamorphic property. To test this idea, we synthesized two peptides: one with the same amino acid sequence as α1 (AYSEKVIDHYENPRN), and the other with an I8K substitution (AYSEKV<u>K</u>DHYENPRN). We analyzed the ^1^H-^15^N HSQC spectra of IscU WT before and after treatment with these peptides and confirmed that the D-state population increased after the addition of the peptides **(Fig. 5 and Extended data Fig. 10)**. Of note, the increase in the D-state population was more noticeable with the I8K peptide treatment compared to the WT-like peptide **(Fig. 5b)**. These results suggest that the α1-like peptide interfered with the interaction between α1 and other regions of IscU, resulting in the loss of constraints imposed by the N-terminus to stabilize the S-state **(Fig. 5a)**. In addition, we observed that the peptide treatment altered the ^1^H-^15^N chemical shifts of the residues in the β1-β3, α2, and α5 motifs of IscU (**Extended data Fig. 10**). These peptide-induced signal perturbations were akin to those observed with the α1-motif variants (**Extended Data Fig. 9**), suggesting that α1-like peptides may induce IscU’s metamorphic alterations through the similar mechanism of the α1-motif variants to modulate their conformational variability.

**Fig 5.**
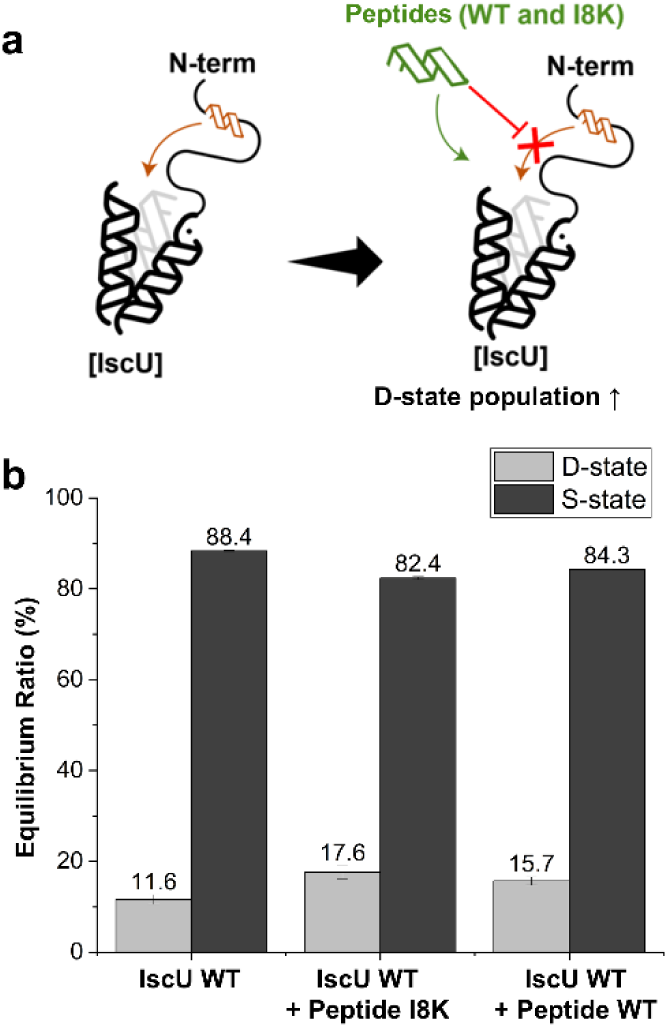
The peptides mimicking the N-terminal α1-helix can change the conformation equilibrium of IscU. **a,** The scheme illustrates the effects of peptides on IscU. **b,** S-state and D-state populations changed by the addition of excess peptides (WT, the peptide of the same sequence with the α1-helix; I8K, the peptide that has the sequence of the α1-helix with the I8K substitution) were measured by monitoring the corresponding signal intensities for K128.

### Role of IscU’s N-terminal Region in Fe-S Cluster Biogenesis

Finally, to verify the importance of the N-terminal region of IscU in the biogenesis of Fe-S clusters, we measured the binding affinity of N-terminal variants of IscU with HscA, an essential Hsp70-like protein in *E. coli* Fe-S cluster biogenesis, using isothermal titration calorimetry (ITC)^22^. Previous studies have reported that HscA interacts with the ^99^LPPVK^103^ motif of IscU, and this interaction stabilizes the D-state of IscU^23–24^. We hypothesized that although the N-terminus of IscU does not directly bind to HscA, the mutations in this region may affect the strength of the interaction. Indeed, we found that the dissociation constant (*K*_d_) for HscA correlated with the α1 stability of IscU variants; I8K showed the weakest affinity, while Y11A and R15A showed increased affinity **(Table 1 and Supplementary Fig. 5)**. This confirmed the significance of the metamorphic equilibrium of IscU in controlling its interaction with HscA. Moreover, it raises an intriguing possibility that the α1 motif could be targeted to modulate the interaction of IscU with other proteins, such as HscA, HscB, IscS, and IscX. Indeed, as observed in the CSP data above **(Extended data Fig. 5 and 9)**, the substitution of Y3 and α1 residues caused NMR signal perturbations in the regions around the LPPVK motif, indicating that the interaction of HscA with this motif may also change the structural state near the N-terminal region.

**Table 1.**
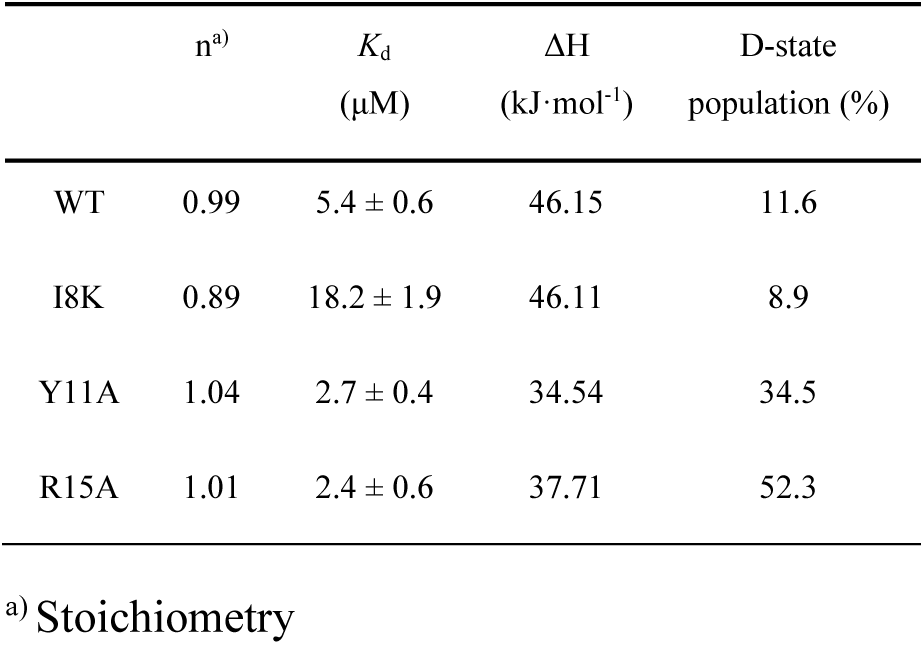
Isothermal titration calorimetry results for the interactions of IscU variants with HscA.

| | n<sup>a)</sup> | $K_d$<br>($\mu\text{M}$) | $\Delta H$<br>($\text{kJ}\cdot\text{mol}^{-1}$) | D-state<br>population (%) |
| --- | --- | --- | --- | --- |
| WT | 0.99 | $5.4 \pm 0.6$ | 46.15 | 11.6 |
| I8K | 0.89 | $18.2 \pm 1.9$ | 46.11 | 8.9 |
| Y11A | 1.04 | $2.7 \pm 0.4$ | 34.54 | 34.5 |
| R15A | 1.01 | $2.4 \pm 0.6$ | 37.71 | 52.3 |

<sup>a)</sup> Stoichiometry

## Discussion

In this study, we established a clear correlation between the metamorphic equilibrium of IscU and the structural heterogeneity in the N-terminal region. Our work provides an important clue to elucidate how IscU maintains its metamorphosis. The α1-visible structural models of IscU show that the α1 and the nearby N-terminal region wrap around the β2, β3, α2, and α4 regions. Therefore, it is not surprising that the stabilization of the intra-interaction involving the N-terminus can fortify the stabilization of the S-state of IscU. However, our data demonstrated that IscU maintains the marginal stability for α1 and heterogeneous intra-interaction at the N-terminus, possibly to confer metamorphic structural propensity on IscU. Our computational and NMR results suggest that the N-terminal region may have direct interactions with the β1, β3, and α3-α5 regions of IscU. Notably, a previous NMR study has reported that residues K89 (at the loop between α3 and α4), N90 (α4), S107 (α5), and E111 (α5) are important for the metamorphic equilibrium^5, 25^. The present study offers a plausible explanation for this; these residues may be, either directly or indirectly, involved in interactions with the N-terminal region.

Heretofore, various proteins have been known to maintain their N-terminus or C-terminus in a disordered state, yet with functional significance^26–27^. The present study shows that the N-terminal tail of IscU also harbors important functionality; its transient stability to form an α-helix and sustain heterogeneous intra-interactions is designed on purpose to maximize its functionality as a versatile and competent mediator in Fe-S cluster biogenesis. We demonstrated that the interaction with HscA depends on the instability of α1, suggesting that the destabilization of α1 may be an efficient strategy to enhance the interaction with HscA and facilitate the process of Fe-S cluster transfer.

Finally, the present study may provide a novel insight into developing an unprecedented therapeutic strategy for the pathogenic mechanisms that are related with abnormal Fe-S cluster biogenesis processes. Previous studies demonstrated that the balance between the S-state and the D-state is fine-tuned to maximize the functionality of IscU, and these structural and functional features of IscU are highly vulnerable to an amino acid substitution^5, 24–25, 28^. For example, the conformational equilibrium could be significantly perturbed by single amino acid substitutions, such as D39A, D39V, K89A, N90A, and E111A for *E. coli* IscU, and D39V and M108I for human ISCU^25, 29^. Notably, it was recently reported that some missense mutations, such as G18E and G64V, can perturb the Fe-S cluster biogenesis mechanism and incur several pathogenic processes in humans^30–31^. Although further studies should be undertaken to appreciate the related molecular mechanisms in detail, it appears likely that these pathogenic mutations alter the conformational balance of IscU, which could be a highly challenging target for therapeutic purposes. Therefore, our novel approach to modulating the equilibrium between the S-state and the D-state by targeting the intra-interaction between the α1 helix and the other parts of IscU may provide unprecedented insight into preventing or suppressing these pathogenic mechanisms. We are conducting additional studies to validate that the α1-like peptide can be a promising therapeutic molecule for abnormal Fe-S cluster biogenesis.

## Methods

### Sequence alignment analysis for the conserved regions of IscU from a few representative species

The species that were selected from the Uniprot database for analyzing the conserved regions of IscU (**Extended data Fig. 7a**) included THEMA (*Thermotoga maritima*; UniProt: Q9X192), STRP1 (*Streptococcus pyogenes* serotype; UniProt: Q9A1G2), AQUAE (*Aquifex aeolicus*; UniProt: O67045), ARCFU (*Archaeoglobus fulgidus*; UniProt: P0DMG1), ARATH (*Arabidopsis thaliana*; UniProt: Q9MAB6), HUMAN (*Homo sapiens*; UniProt: Q9H1K1), MOUSE (*Mus musculus*; UniProt: Q9D7P6), AZOVI (*Azotobacter vinelandii*; UniProt: O31270), ECOLI (*Escherichia coli*; UniProt: P0ACD4), and HAEIN (*Haemophilus influenzae*; UniProt: Q57074)^32^. Alignment of these amino acid sequences was done with Clustal Omega^33^.

### AF-cluster analysis

For AF-cluster analysis, we first generated the multiple sequence alignment (MSA) of IscU. We collected IscU sequences with protein-level, transcription-level, or homology evidence by querying protein names and gene names from the UniProt database. To ensure accurate sequence alignment, we filtered out 1) exceedingly long sequences that contain domains of other proteins, 2) repeatedly reported or duplicated sequences that distort the frequency of each amino acid appearing on the sequence collection, and 3) sequences that were self-reported as low-quality data. With the remaining 6,193 sequences, we ran a Clustal Omega alignment to generate the MSA with each residue corresponding to each other^33–34^. We prepared the full, non-trimmed version of the Clustal Omega MSA for SCA analysis to capture all possible statistical relationships. The trimmed version of the Clustal Omega MSA following the representative *E. coli* sequence (UniProt: P0ACD4) was prepared for AF-cluster analysis to obtain predictions about the intact IscU molecule. Subsequently, the publicly available Colab script version of AF-cluster was imported for its execution^14^. For the input data to run the code, we prepared the trimmed version of the IscU MSA. Under the AF-cluster procedure, the MSA of IscU was clustered by DBSCAN with the eps = 3.5 for the peak number of clusters to optimize between minimal occurrences of outliers and merges^14^. Out of the 260 clusters, 41 clusters had 20 or more sequences as members. These clusters were processed by AlphaFold2 to obtain the corresponding structure predictions^15^. The distances between C_β_ of the N-terminal A2 residue and C_β_ of the others in the AF-cluster models were calculated using Biopython.

### SCA analysis

To execute the core SCA analysis, we imported the MATLAB SCA script, which is available from the original publication^17^. We used non-trimmed MSA of the IscU sequences to capture all possible statistical relationships. Then, spectral cleaning was applied to exclude the statistical and historical noises. Under the spectral cleaning protocol, principal component analysis (PCA) was applied to obtain a simple relationship representation. Out of the total 474 eigenvalues, the lowest 467 eigenvalues were indistinguishable from those of the randomized alignments with the same size and amino acid composition. The corresponding 467 eigenmodes were classified as random noise. The first eigenmode describes the phylogenetic history, thus excluded from the coevolution relationship dataset^16–18^. Scikit-learn was used to execute the *k*-means clustering algorithm to cluster the coevolving residues^35^. The X-ray structural model of IscU (PDB: 3LVL; IscU in the IscU-IscS complex) was used for the structural representation of the protein sectors^8^. Pymol and UCSF ChimeraX were used for visualization and model superposition^36–37^. Microsoft Excel 2019, NumPy, Pandas, and Biopython were used for the data analysis^38–40^. Origin (OriginLab), Matplotlib, and Seaborn were utilized for the graphical data representation^41–42^.

### Preparation of recombinant proteins

We used the pTrc99A-based expression plasmid of IscU WT to incur single-residue substitution via site-directed mutagenesis, for which the site-directed mutagenesis kit (Enzynomics) was employed, and each mutation was verified by DNA sequencing^6^. For protein expression, the expression plasmid was transformed to either *E. coli* BL21[DE3] (for WT, I8K, and R15A) or *E. coli* Rosetta[DE3] (for the other variants) competent cells (Enzynomics). The transformant was grown first overnight on a Luria Broth (miller; LB) plate containing appropriate antibiotics (100 μg/mL ampicillin for BL21[DE3] or 100 μg/mL ampicillin plus 34 μg/mL chloramphenicol for Rosetta[DE3]), and then inoculated and grown sequentially in 3 mL, 100 mL, and 1 L liquid LB media at 37 °C. The cells for the production of uniformly ^15^N-labeled or ^13^C/^15^N-labeled ([U-^15^N] or [U-^13^C, U-^15^N], respectively) protein sample were cultured in 1 L M9 minimal medium, containing 33.7 mM Na_2_HPO_4_, 22 mM KH_2_PO_4_, 8.55 mM NaCl, 9.35 mM NH_4_Cl, 0.4% glucose, 1 mM MgSO_4_, 0.3 mM CaCl_2_, 1 μg biotin, 1 μg thiamin, 134 μM ethylenediaminetetraacetic acid (EDTA), 31 μM FeCl_3_-6H_2_O, 6.2 μM ZnCl_2_, 0.76 μM CuCl_2_-2H_2_O, 0.42 μM CoCl_2_-2H_2_O, 1.62 μM H_3_BO_3_, and 0.081 μM MnCl_2_-4H_2_O. For ^15^N-labeling or ^13^C/^15^N-labeling, ^15^NH_4_Cl and/or ^13^C-glucose (Cambridge isotope laboratories, Inc.) were respectively added to the M9 media instead of their natural-abundance counterparts^43^. Protein expression was induced by treating 1 mM isopropyl β-D-1-thiogalactopyranoside (IPTG) to a cell culture at the OD_600_ of 0.5-0.6. The cells were further grown for 6∼12 hours, and harvested by centrifugation at 6,000 g for 20 minutes at 4 °C. The cell pellets were rinsed with 0.9% NaCl solution, re-centrifuged at 4,000 g for 20 minutes at 4 °C, and then stored at −80 °C. For the purification of IscU, the cells were first resuspended with the buffer, consisting of 50 mM Tris-HCl, 0.5 mM EDTA, and 1 mM dithiothreitol (DTT) at pH 8.0, sonicated for cell lysis, and centrifugated at 40,000 g for 40 minutes at 4 °C to remove cell debris. The supernatant from this was then applied to a HiTrap Q HP 5mL column (Cytiva) with an ÄKTAprime plus fast protein liquid chromatography (FPLC) system (Cytiva), and a NaCl gradient from 0 to 150 mM was taken for IscU elution. We used SDS-PAGE analysis to find the fractions containing IscU. The pooled fractions were concentrated by using a centrifugal concentrator (3 kDa cutoff; Sartorius), and subsequently applied to a Hiload 16/600 Superdex^TM^ 75pg column (Cytiva) that was connected to an ÄKTAgo FPLC system (Cytiva). The buffer for this contained 20 mM Tris-HCl, 0.5 mM EDTA, 150 mM NaCl, and 1 mM DTT at pH 8.0. Again, the fractions containing IscU were selected based on SDS-PAGE analyses, and the final sample was concentrated, aliquoted, flash frozen with liquid nitrogen, and stored at −80 °C until used. For efficient HscA preparation, we attached the His_6_-SUMO tag to the N-terminus of HscA at the original pTrc99A-based HscA expression plasmid^24^. In addition, we produced a recombinant enzyme, His_6_-tagged ubiquitin-like specific protease 1 (the endopeptidase recognizing and cleaving right after the SUMO domain; Ulp1), to cleave out the His_6_-SUMO tag from HscA. Of note, this procedure did not leave any residual amino acid other than the original sequence of HscA. The pET28a(+)-based expression plasmid was used for the overexpression of His_6_-Ulp1. The integrity of these expression plasmids was verified with DNA sequencing. We transformed the *E. coli* BL21[DE3] (for HscA) or Rosetta[DE3] (for Ulp1) competent cells. The transformant was grown first overnight on a LB plate containing appropriate antibiotics, and then inoculated and grown sequentially in 3 mL, 100 mL, and 1 L liquid LB media at 37 °C. To induce overexpression, 0.5 mM IPTG was treated to 1 L LB when the OD_600_ was 0.5-0.6. The cell was further grown for 6-12 hours, and harvested by centrifuging the media at 6,000 g for 20 minutes at 4 °C. The cell pellets were rinsed with the 0.9 % NaCl solution, recollected with centrifugation at 4,000 g for 20 minutes at 4 °C, and stored at −80 °C until used. For purification of HscA and Ulp1, the cell pellets were first resuspended with the buffer, containing 50 mM Tris-HCl, 200 mM NaCl, 5% glycerol, 0.01% Triton X-100, and 20 mM imidazole at pH 7.5, and then lysed by the sonication. The supernatant was retrieved by centrifuging the lysate at 40,000 g for 40 minutes at 4 °C, and applied to a Histrap HP 5 mL column (Cytiva) on ÄKTAprime plus for purification. The sample containing the target protein was eluted by the elution buffer consisting of 20 mM Tris-HCl, 200 mM NaCl, and 500 mM imidazole at pH 7.5. The fractions were analyzed by SDS-PAGE to identify the target protein and estimate its purity. At this stage, no further purification was necessary for Ulp1, whereas HscA required additional cleavage and purification procedures. First, to remove the tag, the His_6_-SUMO-HscA sample was treated with Ulp1 in a 1:200 molar ratio. The mixture was dialyzed at room temperature overnight by using SnakeSkin^TM^ Dialysis Tubing (10 kDa cut off; ThermoFisher). The dialysis buffer contained 20 mM Tris-HCl and 100 mM NaCl at pH 7.5. After SUMO cleavage, the sample was again applied to a Histrap HP 5 mL column, at which tag-removed HscA did not bind to a column and was eluted as a flow-through. The pooled fraction was finally applied to a Hiload 16/600 superdex^TM^ 75 pg column with the elution buffer of 50 mM Tris-HCl, 150 mM NaCl, 0.5 mM EDTA, and 1 mM DTT at pH 7.5. The cleavage and purification of HscA were monitored with SDS-PAGE. The purified fractions were concentrated, aliquoted, flash frozen with liquid nitrogen, and stored at −80 °C until used.

### Circular dichroism (CD) spectroscopy

The far-UV CD spectra at the wavelength range of 190 to 260 nm were measured by using the JASCO J-1500 CD spectrophotometer at 25 °C. The concentration of IscU samples was 20 μM in the buffer containing 20 mM Tris-HCl, 150 mM NaCl, and 5 mM tris(2-carboxyethyl) phosphine hydrochloride (TCEP) at pH 8.0. The cuvette with the pathlength of 1 mm was used. For the temperature titration experiments from 15 to 75 °C, the CD signal at 222 nm was measured at every 5 °C increase with a 5-min incubation period. All data sets were analyzed with the Spectra Manager software (JASCO). Graph fitting and transition temperature calculation were performed with the Boltzmann model in Origin^44^.

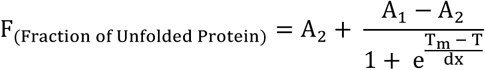

where A_2_ represents the temperature at which the protein is fully unfolded, while A_1_ corresponds to temperature at which it is fully folded. T_m_ denotes the melting temperature (midpoint between A_1_ and A_2_), and T is the experimental temperature. The α-helix composition was calculated from the CD dataset using the Bestsel website^45^.

### Size exclusion chromatography (SEC)

The SEC analysis was performed using a Superdex™ 75 Increase 10/300 GL column (Cytiva) on an ÄKTA pure (Cytiva). The buffer for this analysis consisted of 20 mM Tris-HCl, 0.5 mM EDTA, 150 mM NaCl, and 1 mM DTT at pH 8.0. The concentration of all the protein samples was 0.1 mM. The 280 nm absorbance chromatograms were normalized by Origin.

### Nuclear magnetic resonance (NMR) spectroscopy

All NMR data were acquired with the Advance III HD 850 MHz NMR spectrometer equipped with a cryogenic HCN probe (Bruker). The buffer for the NMR samples was composed of 20 mM Tris-HCl, 0.5 mM EDTA, 150 mM NaCl, 5 mM DTT, 7% D2O, and 1.5 mM 3-(trimethylsilyl)-1-propane sulfonic acid sodium salt at pH 8.0. Samples were filled into 5 mm Shigemi tubes (SHIGEMI Co.) with a volume of 0.3 mL. For 2D ^1^H-^15^N HSQC spectra, the [U-^15^N]-IscU WT and variant samples were prepared at a concentration of 0.1 mM. For backbone signal assignments, the [U-^13^C, U-^15^N]-IscU variant samples were prepared at 0.7 mM, and the following spectral data were collected: 2D ^1^H-^15^N HSQC, 3D HNCA, 3D HN(CO)CA, 3D HNCO, 3D HN(CA)CO, 3D HNCACB, and 3D CBCA(CO)NH. NMR data acquisition and processing were performed with Topspin 3.2 (Bruker), while all NMR data analyses were conducted with POKY software^46^. The chemical shift perturbation (CSP) between IscU WT and its variants was calculated with the following equation.

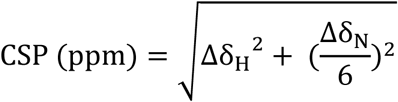

where Δδ_H_ and Δδ_N_ indicate differences in ^1^H and ^15^N chemical shifts between WT and variants, respectively. The combined secondary ^13^C chemical shift for IscU WT and its variants was calculated with the following equation^47–48^.

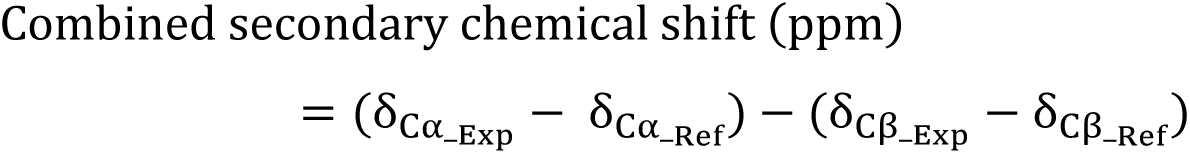

where δ_Cα_Ref_ and δ_Cβ_Ref_ respectively denote the referencing chemical shifts of C_α_ and C_β_, while δ_Cα_Exp_ and δ_Cβ_Exp_ denote the experimental chemical shifts of C_α_ and C_β_, respectively.

The conformational equilibrium ratios (S-state vs. D-state) for IscU variants were analyzed by comparing the intensities of the K128 peaks corresponding to the S- and D-states in the ^1^H-^15^N HSQC spectrum (**Supplementary Fig. 2**). The errors in these analyses were estimated based on the peak intensity/signal-to-noise ratio. The secondary structure predictions were made by feeding the chemical shift values of C_α_, C_β_, and H_N_ into a TALOS-N software^49^. For the measurement of ^1^H-^15^N heteronuclear nuclear Overhauser effects (NOE), we used the sample containing the IscU variant with a concentration of 0.5 ∼ 0.7 mM. Data analysis was performed by using the analysis script implemented in the POKY software package^46^.

### PRE experiment

For PRE measurement, we used a stable nitroxide radical, 4-maleimido-TEMPO (Sigma-Aldrich), which can target a free thiol group of a cysteine residue through a thiol-maleimide reaction^20^. First, we exchanged the buffer of the [U-^15^N]-IscU A2C sample with a buffer containing 10 mM EDTA, 20 mM Tris-HCl, 150 mM NaCl, and 1 mM DTT at pH 8.0 to remove any metal ions. Subsequently, we exchanged the buffer to the one containing 20 mM Tris-HCl and 150 mM NaCl at pH 7.0, and treated the 2-fold excess of zinc chloride to the sample. This step was essential to protect the native cysteine residues, which coordinate the zinc ion and become resistant to cysteine-modifying chemicals. We also used the pH 7.0 buffer to avoid a thiol-maleimide reaction at an amine group of lysine^20^. After a 30-minute incubation, the sample containing zinc-bound IscU A2C was treated with 0.75 equivalents of 4-maleimido-TEMPO for 10 minutes at room temperature. In this step, the concentration and incubation time for 4-maleimido-TEMPO treatment were optimized to minimize any unwanted reaction. The sample was then washed with a buffer containing 20 mM Tris-HCl and 150 mM NaCl at pH 8.0 to remove unreacted chemicals, and 6 mM EDTA was re-added to chelate zinc ions from IscU. The final buffer for NMR measurement was the same with the NMR buffer stated above. The first 2D ^1^H-^15^N HSQC spectrum was acquired with this sample. Subsequently, the sample was treated with excess DTT (5 mM) to reduce a nitroxide radical, and the second 2D ^1^H-^15^N HSQC spectrum was acquired for a reduced sample. The PRE for each residue was calculated by dividing the signal intensity of the first spectrum (a radical-containing sample) with the one of the second spectrum (a reduced sample). To verify the attachment of TEMPO on IscU A2C, the UV-visible spectra (250-500 nm) were obtained for the samples of TEMPO- treated A2C, TEMPO-treated WT, and TEMPO-untreated A2C (**Extended data Fig. 3d-e**). We observed that the TEMPO-treated A2C sample showed the characteristic UV-Vis signal of an oxidized TEMPO moiety, while other samples did not show a similar signal, indicating that our procedure was successful to attach a nitroxide radical at the C2 position of the A2C construct (**Extended data Fig. 3e**)^50^.

### Peptide titration experiment

Two α1-like peptides, mimicking the amino acid sequence of the α1 helix, were synthesized with the following sequences: the WT-like peptide (^2^AYSEKVIDHYENPRN^15^) and the I8K-like peptide (^2^AYSEKV<u>K</u>DHYENPRN^15^). The peptides were directly dissolved in the NMR buffer, consisting of 20 mM Tris-HCl, 150 mM NaCl, 0.5 mM EDTA, and 5 mM DTT at pH 8.0, which was subsequently added to the sample of 0.1 mM [U-^15^N]-IscU WT. The final concentration of the peptides was 0.9 mM. The ^1^H-^15^N HSQC spectra were measured before and after the addition of peptides, and the ratio between S-state and D-state was calculated by comparing the intensities of the corresponding signals of K128.

### Isothermal titration calorimetry (ITC)

We employed a nano ITC instrument (TA instruments) to investigate the binding interaction between IscU mutants and HscA. All samples were first buffer-exchanged to the buffer containing 20 mM Tris-HCl, 150 mM NaCl, 0.5 mM EDTA, and 5 mM TCEP at pH 8.0. IscU were loaded into a syringe in a concentration of 1.5 mM, and HscA was loaded into a sample cell in a concentration of 0.15 mM. ITC experiments, consisting of 25 injections of IscU samples into HscA, were conducted at 25 °C. Data acquisition, processing, and analysis were performed using Nanoanalyze software (TA instruments).

### Size-exclusion chromatography with multi-angle light scattering (SEC-MALS)

We employed the HPLC-MALS instrument (Waters^TM^) to verify the monomeric state of IscU Y11P. A 5 mg/mL sample of IscU Y11P was injected into a BioSuite^TM^ 125 4 μm UHR SEC 4.6 mm x 30 cm column. The buffer for IscU Y11P consisted of 20 mM Tris-HCl, 150 mM NaCl, and 5 mM DTT at pH 8.0. The molecular weight of IscU Y11P was calculated by comparing scattering intensity values from a bovine serum albumin, a well-characterized reference protein.

---

**Extended Data Fig 1.**
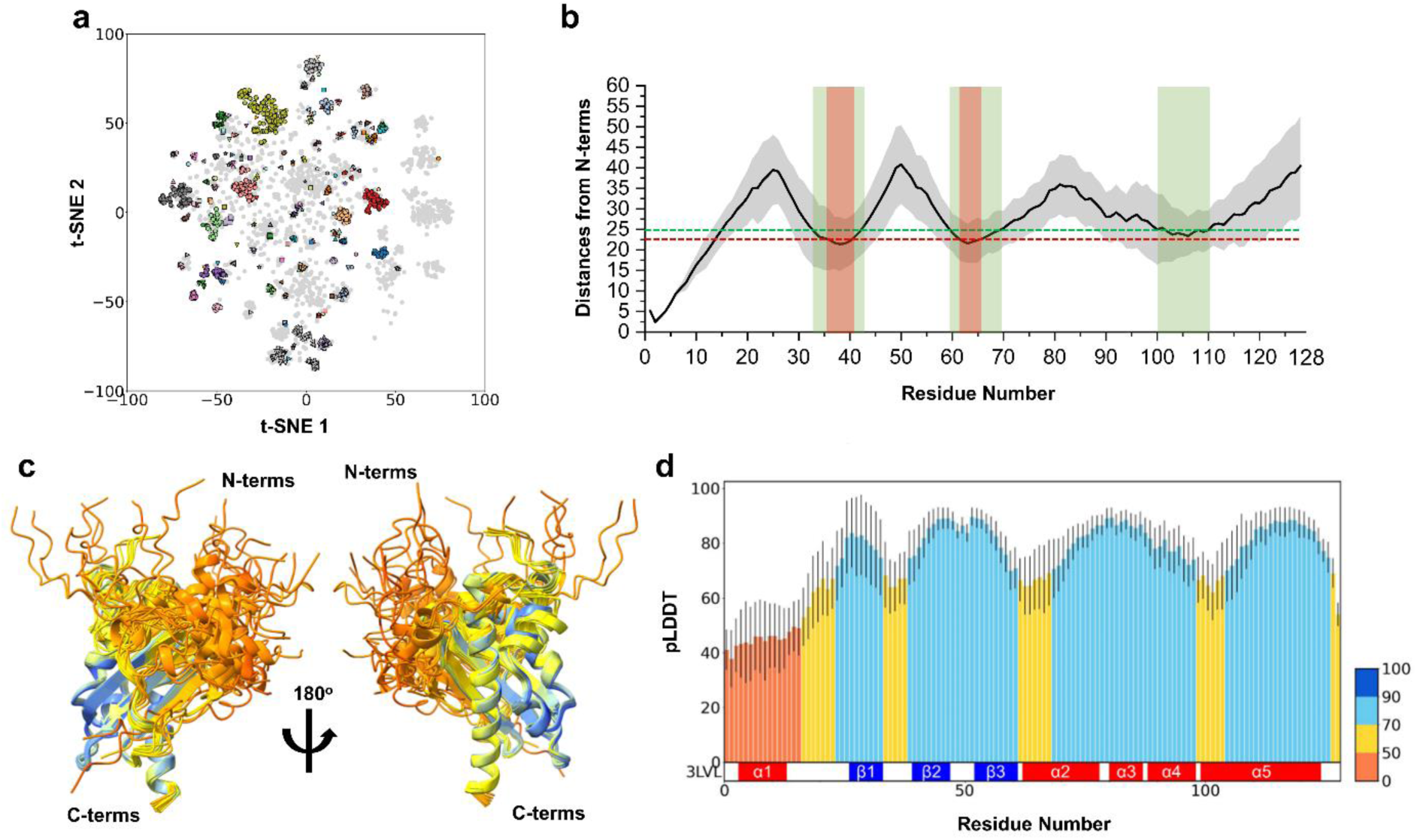
The pLDDT score per residue of the structural models predicted by AF-cluster. **a,** AF-cluster method showed the various clusters in the multiple sequence analysis (MSA) of IscU, by the DBSCAN with the eps=3.5. **b**, Calculated distances for each residue from the N-terminal residue (A2) of AF-cluster models of IscU, along with their standard deviations. The red dashed line and highlighted regions indicate the residues whose distances from A2 are below 23 Å, and the green dashed line and highlighted regions indicate the residues whose distances from A2 are below 25 Å^21^. **c,** Superposition of the AlphaFold2 (AF2) predictions of the IscU MSA clusters with more than 20 sequences each, following the AF2 coloring scheme based on the pLDDT score. A low pLDDT score (orange: < 50, yellow: 50∼70) implies an uncertain per-residue prediction of the structure, and the corresponding motif has a higher possibility of being disordered^14–15^. **d,** The bar plot of the averaged pLDDT score per residue with a standard deviation (shown as a black line). Note that the N-terminal region, especially α1 motif, is classified as a low-confidence prediction region (pLDDT < 50) under the AlphaFold criteria^15^.

**Extended Data Fig 2.**
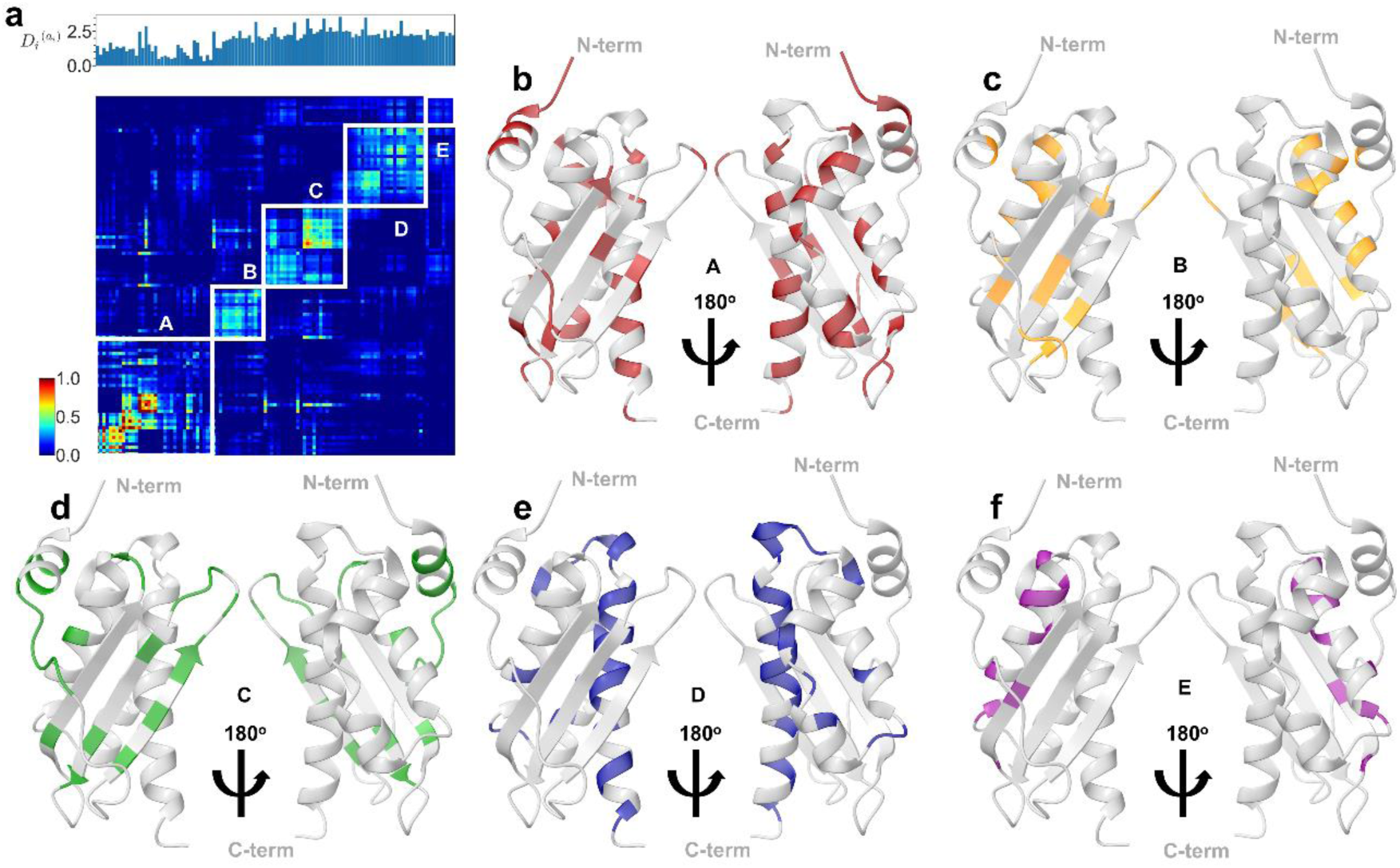
The SCA ‘sectors’ of IscU residues that are clustered by their coevolutionary relationships. **a,** The statistical coupling analysis (SCA) of IscU (Fig. 1c) indicated that IscU’s residues are clustered into 5 ‘sectors’ (denoted as A to E boxes) of the highly coevolving IscU residues by *k*-means^16–18^. Each sector is composed with the following residues: **b**, Sector A: the residues 1, 2, 3, 4, 5, 8, 12, 19, 20, 31, 36, 42, 46, 48, 51, 52, 54, 61, 73, 77, 82, 83, 84, 89, 91, 94, 95, 98, 104, 108, 115, 116, 118, 119, 122, 125, (**c**) Sector B: the residues 11, 21, 22, 27, 29, 34, 40, 43, 44, 56, 65, 69, 72, 76, 93, 96, (**d**) Sector C: the residues 6, 7, 9, 10, 13, 14, 15, 16, 17, 18, 28, 30, 32, 33, 35, 37, 38, 39, 41, 45, 47, 55, 62, 63, 71, (**e**) Sector D: the residues 74, 79, 80, 85, 88, 90, 97, 99, 100, 101, 102, 103, 105, 106, 107, 109, 110, 111, 112, 113, 114, 117, 120, 121, 124 and (**f**) Sector E: the residues 53, 57, 64, 66, 67, 68, 70, 75, 78. Of note, this result clearly shows that residues in the N-terminal region coevolve with other residues of IscU. The X-ray structural model of IscU (PDB: 3LVL) was used for representation of the protein sectors^8^.

**Extended Data Fig 3.**
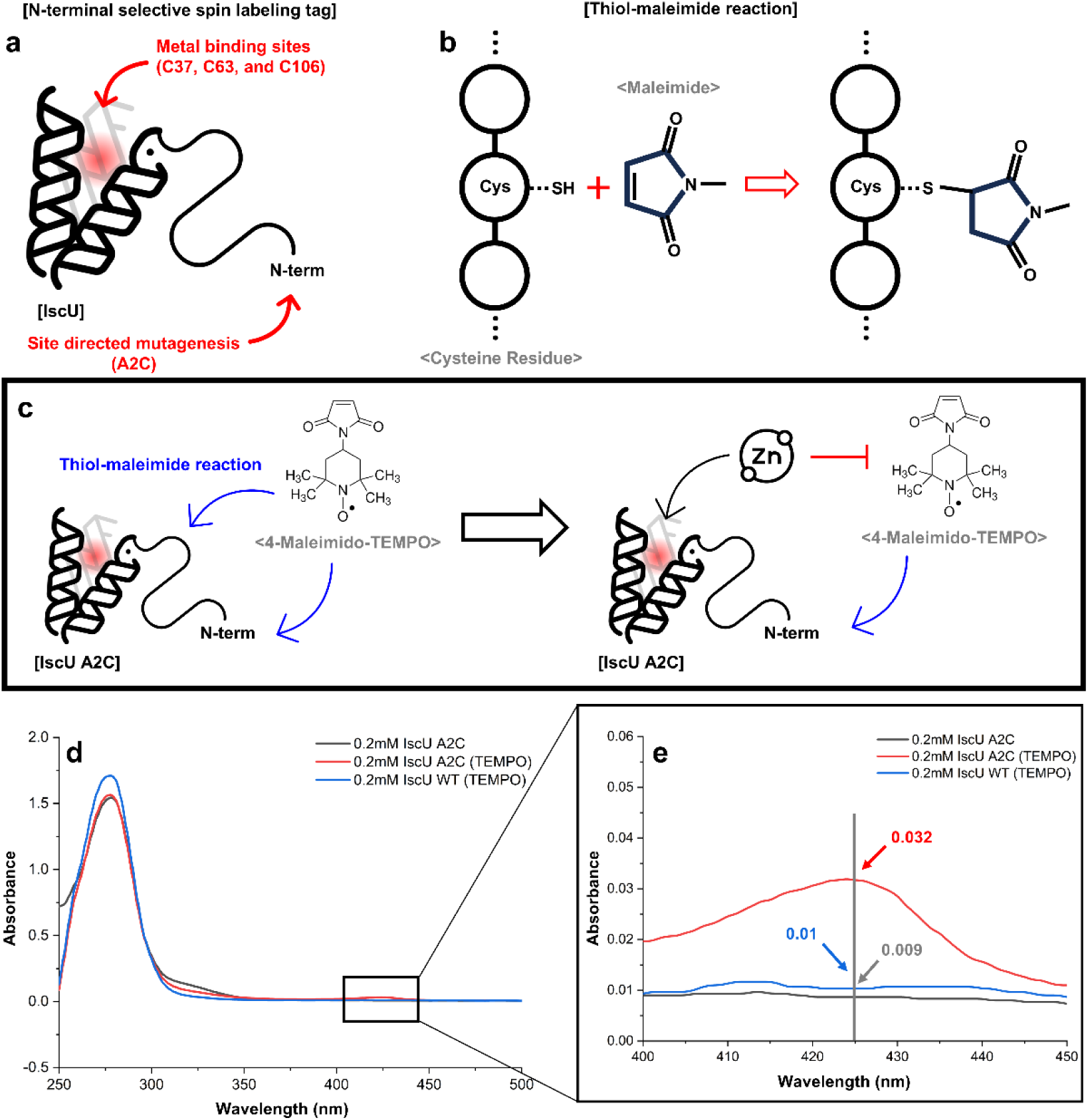
The procedure for the paramagnetic relaxation enhancement (PRE) experiment on IscU. **a-c,** The protocol is to attach a nitroxide radical (4-maleimido-TEMPO) selectively at the C2 position of the IscU A2C variant. We made the A2C variant to covalently attach a nitroxide radical at its C2 position (**a**) by employing 4-maleimido-TEMPO and inducing a thiol-maleimide reaction (**b**). However, IscU has three native cysteine residues (C37, C63, and C106, constructing the metal-binding site; highlighted by a red sphere), which could be targeted by 4-maleimido-TEMPO as well^20^. **c,** To induce the attachment of a radical only at C2 and block the reaction with the other cysteines, we first treated excess zinc chloride, because the native cysteine residues would coordinate zinc ions and exhibit a reduced reactivity toward 4-maleimido-TEMPO. After the reaction, the samples were washed to remove unreacted chemicals, and then excess EDTA was treated to take out a zinc ion from a protein. **d-e,** The UV-Vis spectroscopy measurement to confirm an attachment of a nitroxide radical on IscU A2C. **e,** UV-Vis spectra (250 – 500 nm) of untreated IscU A2C (grey), TEMPO-treated IscU A2C (red), and TEMPO- treated IscU WT (blue) are shown. **e,** Magnified UV-Vis spectra at 400 – 450 nm. The signal at around 425 nm indicated the presence of oxidized TEMPO moiety, which was produced after air exposure of the TEMPO-treated A2C sample for 1-2 days. The same procedure failed to give a similar signal for an untreated A2C or TEMPO- treated WT sample.

**Extended Data Fig 4.**
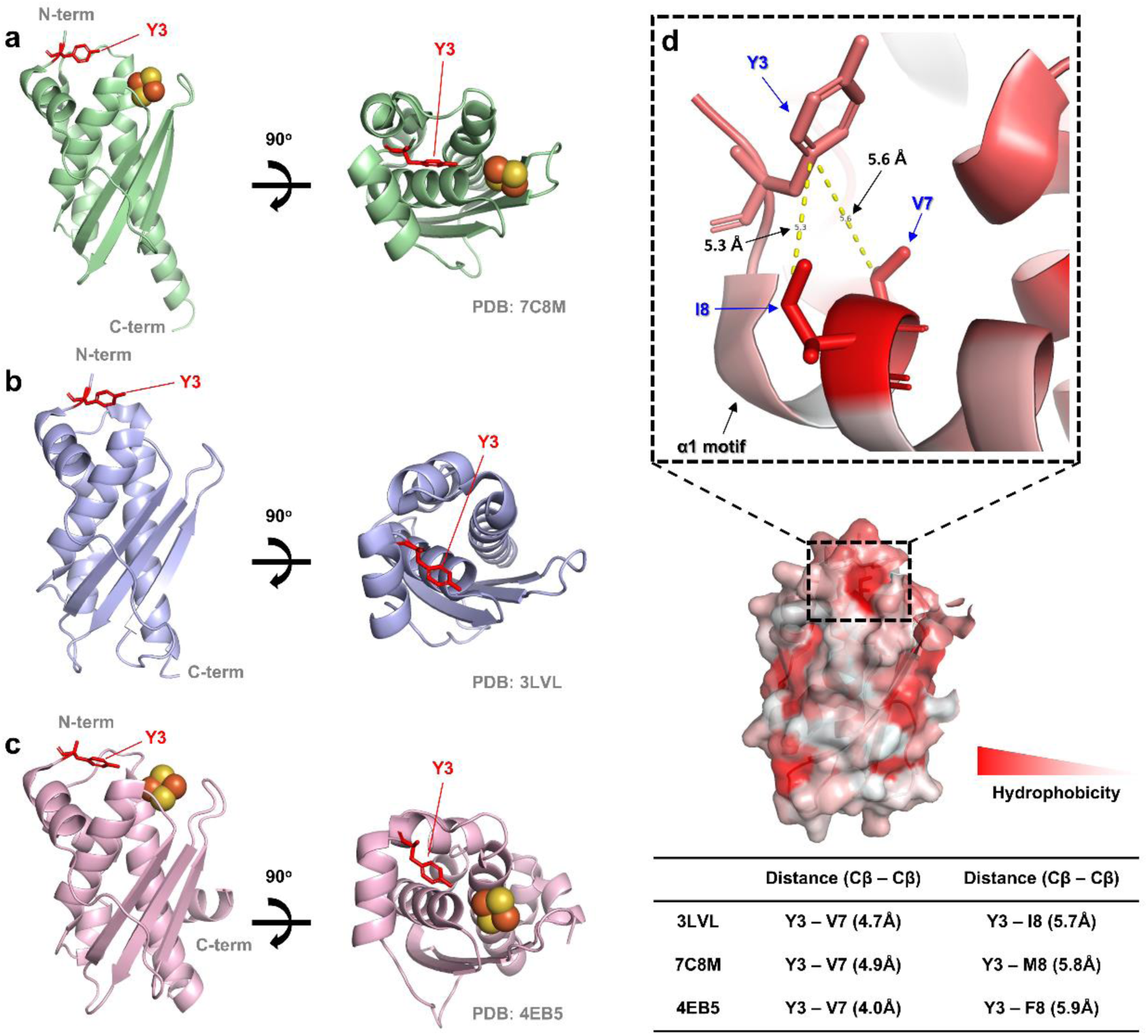
The side chain of Y3 faces toward the core of IscU. The side chain of the highly conserved Y3 residue is shown red in three representative structural models of IscU. **a**, The structural model of IscU from *Methanothrix thermoacetophila* (PDB 7C8M)^9^. **b**, The structural model of IscU from the IscU-IscS complex in *E. coli* (PDB 3LVL)^8^. **c**, The structural model of IscU from the IscU-IscS complex in *A. fulgidus* (PDB 4EB5)^10^. **d**, The red color intensity corresponds to the degree of hydrophobicity. The side chain of the Y3 residue is oriented toward the hydrophobic residues of α1. The C_β_ distances from Y3 to the two hydrophobic residues of α1, V7 and M8, range from 4.0 to 4.9 Å and 5.7 to 5.9 Å, respectively, suggesting hydrophobic packing in this region^51^.

**Extended Data Fig 5.**
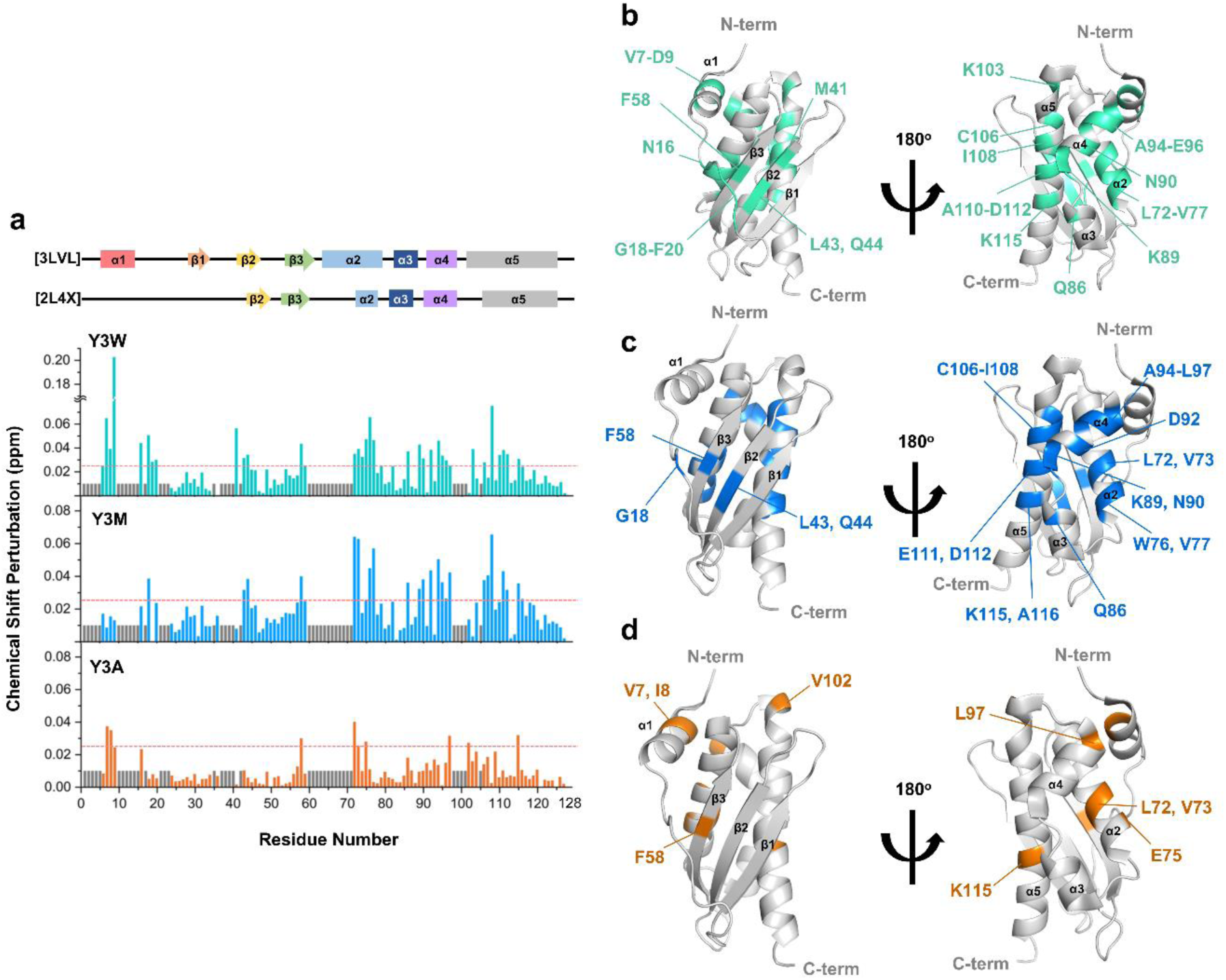
The chemical shift perturbation (CSP) profiles of IscU by Y3 substitutions. **a**, The CSP plots showing the signal shifts from the ^1^H-^15^N HSQC spectra of WT to the same spectra of Y3 mutants (teal blue: Y3W, sky blue: Y3M, orange: Y3A). Unassigned residues are denoted by short grey bars, and the red dashed line represents an arbitrary cut-off to mark the residues exhibiting significant perturbations. The secondary structure from the structural models of IscU WT (PDB 3LVL & 2L4X) is displayed on the top. **b-d**, The residues showing CSP more than the red dashed line in the CSP plot are colored on the structural model of IscU (teal blue: Y3W, sky blue: Y3M, orange: Y3A; PDB 3LVL).

**Extended Data Fig 6.**
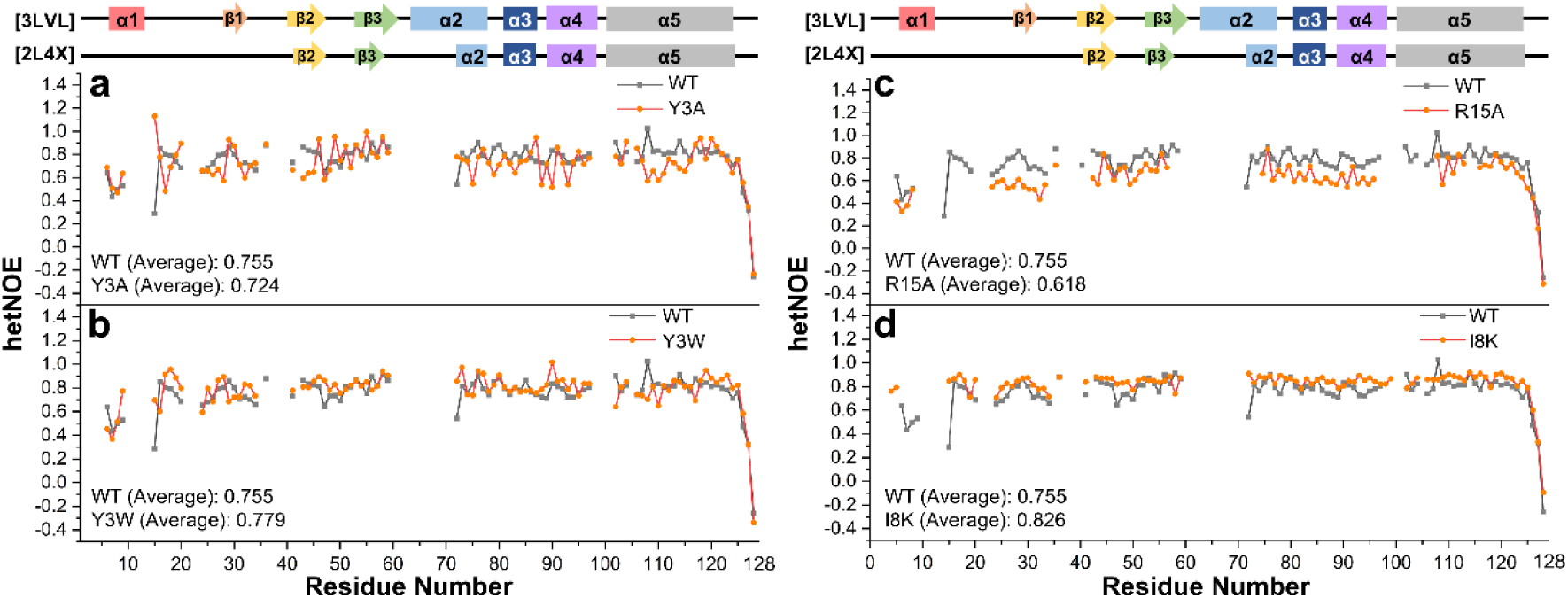
The ^1^H-^15^N heteronuclear NOE (het-NOE) measurements of IscU WT and mutants. **a-d**, The ^1^H-^15^N het-NOE results of IscU mutants (Y3A [**a**], Y3W [**b**], R15A [**c**], and I8K [**d**]; orange) are shown with the results of WT (grey). The secondary structures from the structural models of IscU WT (PDB 3LVL & 2L4X) are displayed on the top.

**Extended Data Fig 7.**
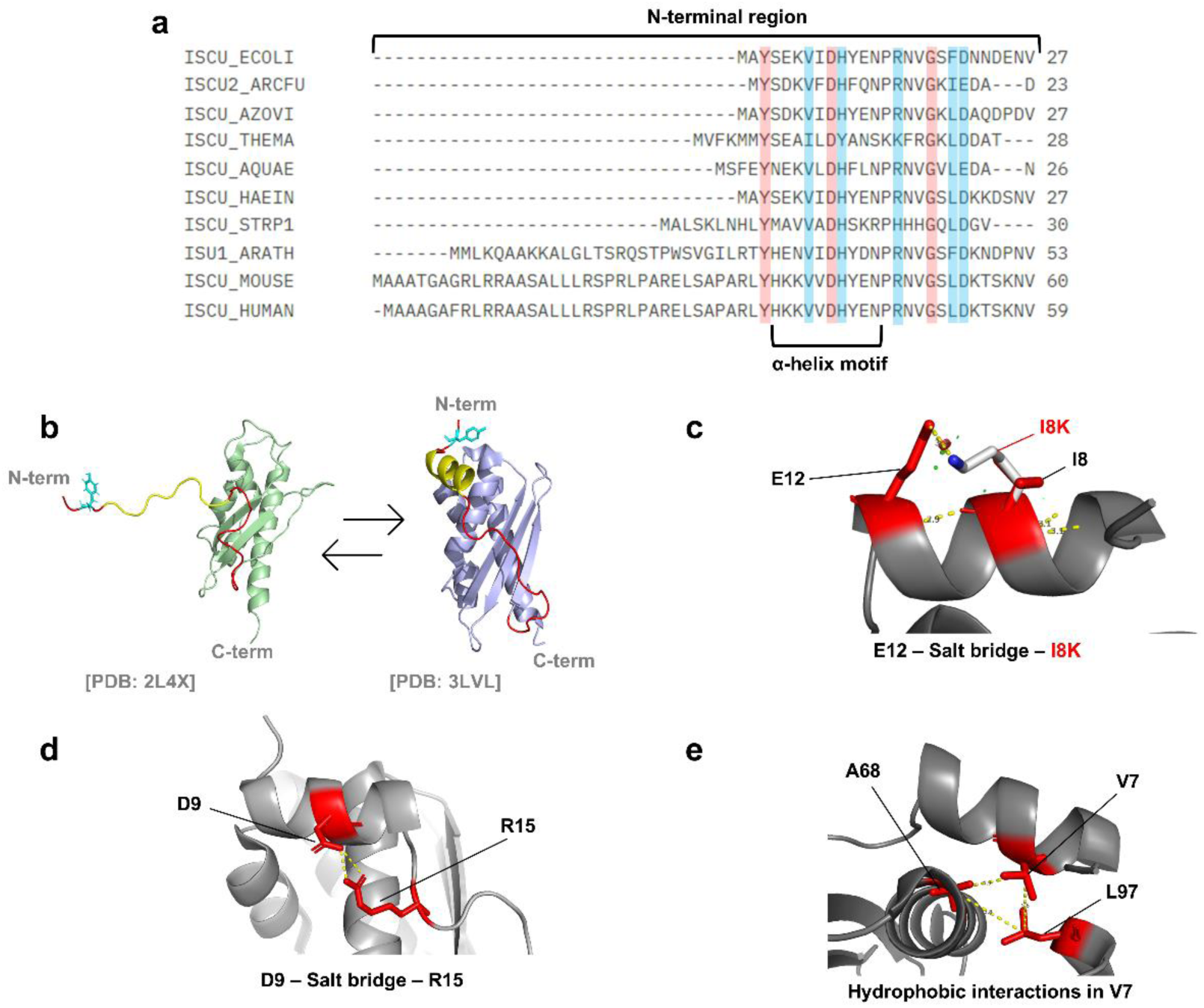
The rationales of the site-directed mutagenesis design for the α1-helix and the nearby regions of IscU. **a**, The amino acid sequence alignment results for the N-terminal region of IscU from a few representative species (ECOLI: *Escherichia coli*, ARCFU: *Archaeoglobus fulgidus*, AZOVI: *Azotobacter vinelandii*, THEMA: *Thermotoga maritima*, AQUAE: *Aquifex aeolicus*, HAEIN: *Haemophilus influenzae*, ARATH: *Arabidopsis thaliana*, MOUSE: *Mus musculus*, HUMAN: *Homo sapiens*) conducted with Clustal Omega^33^. The conserved residues are color-shaded; red indicates the identical residues, and blue indicates the conserved residues. The α1-helix motif (S4-N13 in *E. coli*) is marked below. **b**, Two structural models (PDB 2L4X [left] & 3LVL [right]) showcasing the N-terminal region of IscU. The N-terminal region and the α1-helix region are shown red and yellow respectively. Note that the stable α1-helix was not observed in the solution NMR model (left). The Y3 residue is colored cyan. **c-e**, Intra-molecular interactions within the N-terminal region as observed from the PDB model 3LVL. **c**, The positively charged side chain of K8 in the I8K mutant of IscU is predicted to form a salt bridge with the negatively charged side chain of E12. **d**, D9 makes an ionic interaction with R15. **e**, V7 makes hydrophobic interactions with L97 and A68.

**Extended Data Fig 8.**
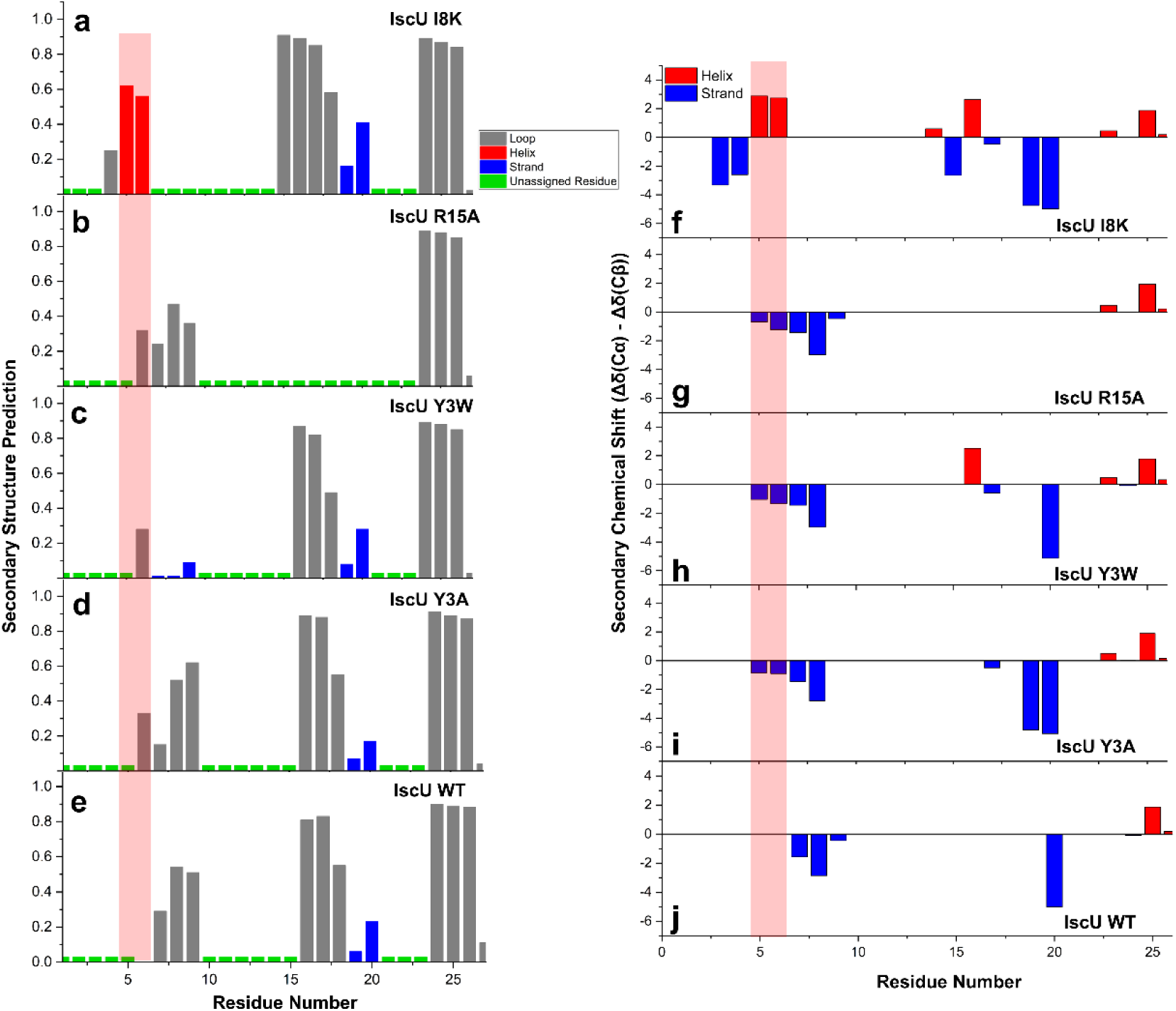
The increased propensity of α-helix formation by I8K mutation. **a-e**, The secondary structure prediction results of TALOS-N for IscU mutants^49^. The following color codes designate the prediction results: red – helix, blue – strand, grey – loop. Unassigned residues are marked with small green squares. Note that TALOS-N calculated an increased propensity of α1-helix formation only in I8K (red shade). **f-j**, The combined secondary chemical shift analysis of IscU mutants based on their C_α_ and C_β_ chemical shifts^47–48^. Red denotes the positive value, suggesting the presence of an α-helix, while blue denotes the negative value, suggesting an β-strand. The bigger value indicates the higher propensity to form a secondary structure. Only the I8K mutant exhibited the positive value at E5 and K6.

**Extended Data Fig 9.**
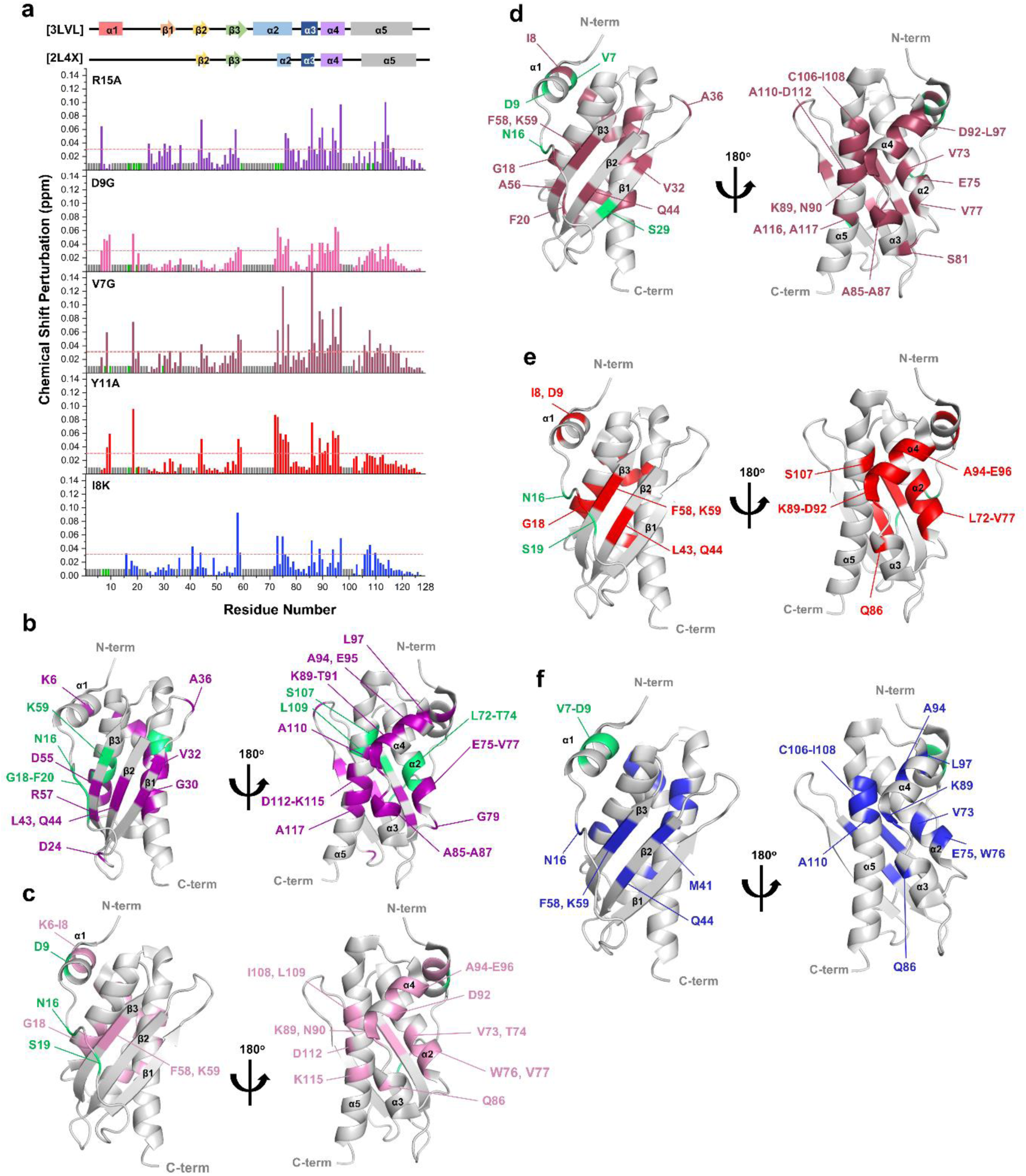
The chemical shift perturbation (CSP) profiles of IscU by α1-helix mutations. **a**, The CSP plots showing the signal shifts from the ^1^H-^15^N HSQC spectra of WT to the same spectra of α1-helix mutants (purple: R15A, pink: D9G, dark red: V7G, red: Y11A, and blue: I8K). The residues, whose signals were visible in WT, but became unassignable in α1-helix mutants, are denoted by green bars. Unassigned residues are denoted by short grey bars, and the red dashed line represents an arbitrary cut-off to mark the residues exhibiting significant perturbations. The secondary structure from the structural models of IscU WT (PDB 3LVL & 2L4X) is displayed on the top. **b-d**, The residues showing CSP more than the red dashed line in the CSP plot are colored on the structural model of IscU (purple: R15A [**b**], pink: D9G [**c**], dark red: V7G [**d**], red: Y11A [**e**], and blue: I8K [**f**]; PDB 3LVL). The residues whose signals disappeared in α1-helix mutants are colored green.

**Extended Data Fig 10.**
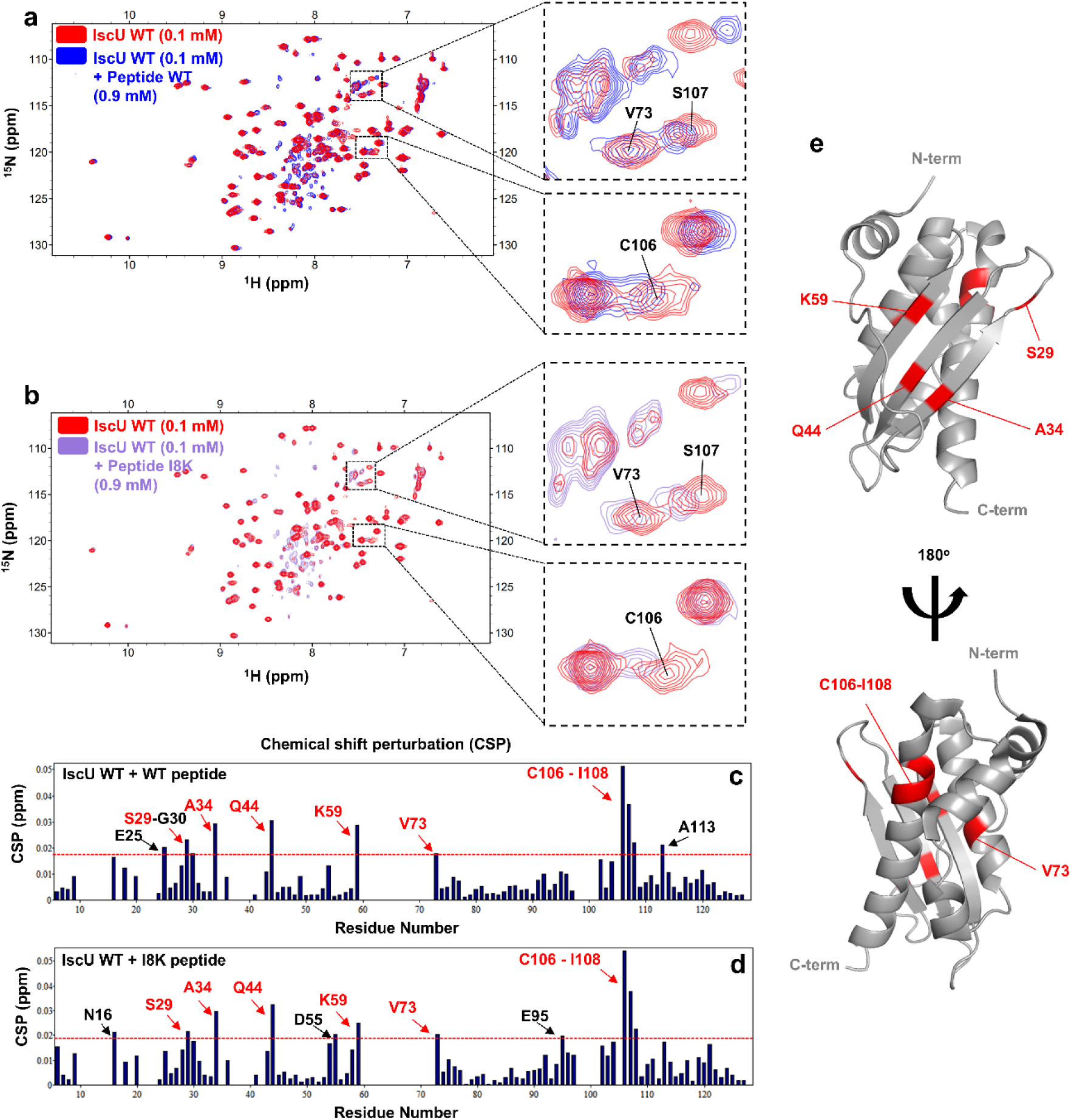
The spectral change of IscU WT by the addition of α1-like peptides. **a**, The ^1^H-^15^N HSQC spectra of 0.1 mM IscU WT taken before (red) and after (blue) the addition of 0.9 mM α1-like I8K peptide are overlaid. **b**, The ^1^H-^15^N HSQC spectra of 0.1 mM IscU WT taken before (red) and after (purple) the addition of 0.9 mM α1-like peptide are overlaid. **c-d**, The chemical shift perturbation (CSP) plots showing the weighted perturbations of the ^1^H-^15^N HSQC signals of IscU upon comparing them to the signals of IscU titrated with α1-mimic peptides ([**c**] WT peptide; [**d**] I8K peptide). The dashed red line represents an arbitrary cut-off to mark the residues exhibiting significant perturbations. Red-colored residues showed significant CSPs in both the WT and I8K peptide titrations. **e**, The red-colored residues in the panels (**c**) and (**d**) are highlighted on the structural model of IscU (PDB 3LVL).

## Acknowledgements

The authors gratefully acknowledge the use of the NMR facility at Pohang University of Science and Technology, Republic of Korea. This work was supported by the National Research Foundation of Korea (NRF) grant [NRF-2020R1I1A2074335 (J. H. K.); NRF-2023R1A2C1006248 (W. Y.)], the Institute for Basic Science of Korea [IBS-R014-A1 (Y. H. K.)], and the TÜBİTAK(Türkiye)-NRF(Korea) 2516 bilateral research program [project number 221N005 (H. D.) and NRF-2021K2A9A1A06096295 (J. H. K.)].

## Notes

### Competing Interest Statement

The authors have declared no competing interest.

