## Supplementary Figures for "The N-terminal order-disorder transition is a critical determinant for a metamorphosis of IscU"

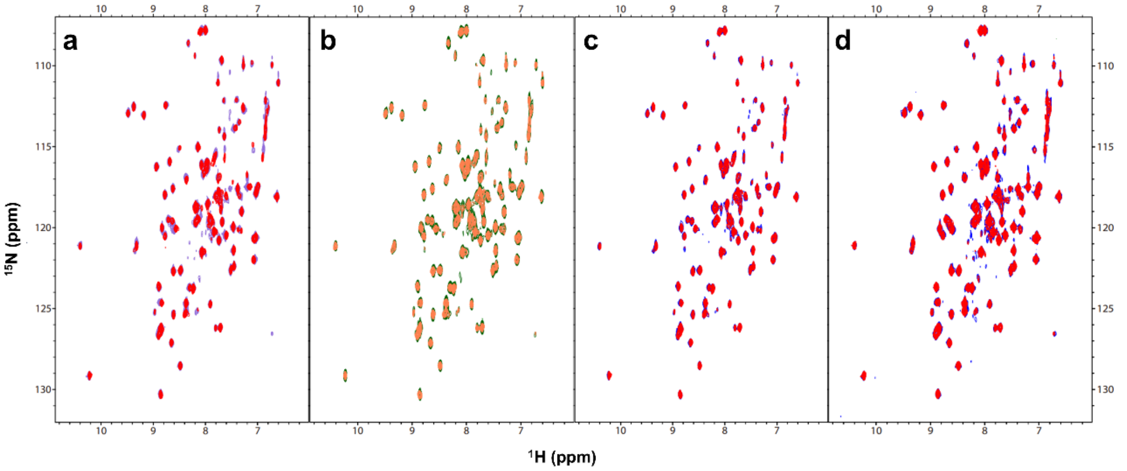


**Supplementary Fig 1. The $^{1}$H-$^{15}$N HSQC spectra of 0.2 mM IscU WT and A2C before and after the attachment of a nitroxide radical (TEMPO)**

**a**, The overlapped $^{1}$H-$^{15}$N HSQC spectra of TEMPO-treated 0.2 mM WT (purple) and TEMPO-treated A2C (red). **b**, The overlapped $^{1}$H-$^{15}$N HSQC spectra of untreated (green) and TEMPO-treated A2C (orange). Overall spectral similarity indicates that neither A2C substitution nor TEMPO attachment caused significant structural perturbation. **c**, The overlapped $^{1}$H-$^{15}$N HSQC spectra of TEMPO-treated A2C before (red) and after (blue) adding DTT. **d**, The overlapped $^{1}$H-$^{15}$N HSQC spectra of TEMPO-treated WT before (red) and after (blue) adding DTT.

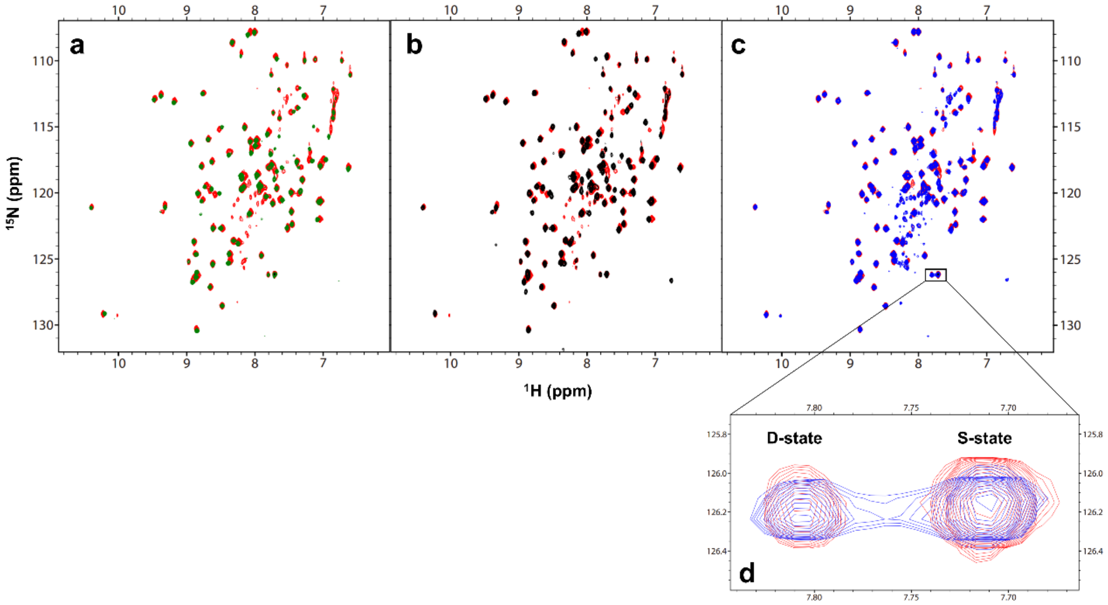


**Supplemenatry Fig 2. The overlapped $^{1}$H-$^{15}$N HSQC spectra of IscU Y3 mutants**

**a**, The $^{1}$H-$^{15}$N HSQC spectrum of 0.1 mM IscU Y3W (green) overlapped with the same spectra of 0.1 mM IscU WT (red). **b**, The $^{1}$H-$^{15}$N HSQC spectrum of 0.1 mM IscU Y3M (black) overlapped with the same spectra of 0.1 mM IscU WT (red). **c**, The $^{1}$H-$^{15}$N HSQC spectrum of 0.1 mM IscU Y3A (blue) overlapped with the same spectra of 0.1 mM IscU WT (red). **d**, The magnified view of the K128 signals in the overlapped $^{1}$H-$^{15}$N HSQC spectra of IscU WT (red) and Y3A (blue). K128 exhibits two concomitant peaks; the left and right peaks represent the D-state and S-state, respectively. The normalized intensity of these two peaks can be used to estimate the conformational equilibrium ratio between the two states[1].

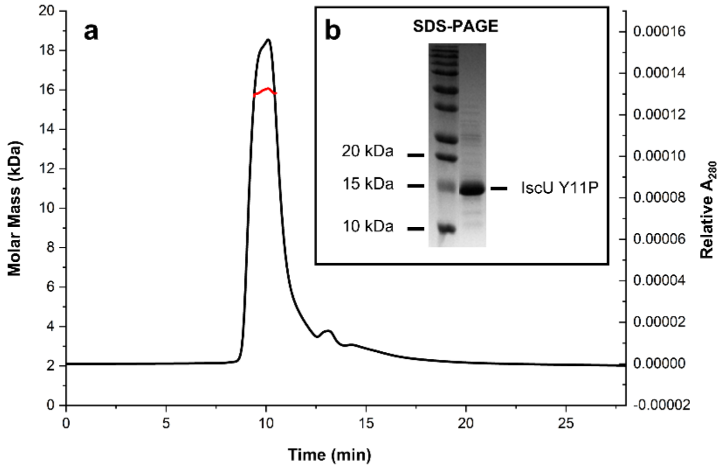


**Supplementary Fig 3. Size exclusion chromatography with multi-angle light scattering (SEC-MALS) analysis of IscU Y11P**

**a**, The SEC-MALS analysis of IscU Y11P verified that its molecular weight was approximately 15-16 kDa. The black line represents the SEC profile of Y11P, while the red line denotes the molar mass of the corresponding SEC peak. **b**, The SDS-PAGE analysis of the IscU Y11P sample showed that the molecular weight of IscU Y11P was approximately 15 kDa (Lane 1: protein marker; Lane 2: IscU Y11P).

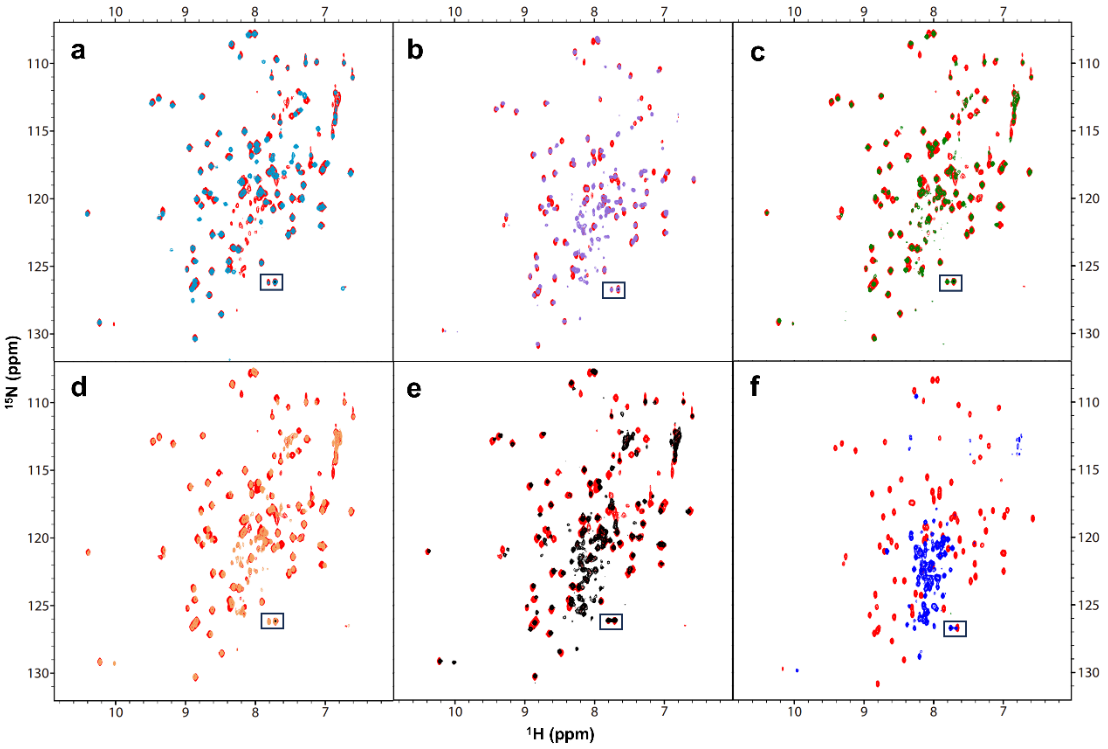


**Supplementary Fig 4. The overlapped $^{1}$H-$^{15}$N HSQC spectra of IscU α1-helix mutants**

The $^{1}$H-$^{15}$N HSQC spectra of 0.1 mM IscU α1-helix mutants (sky blue: I8K [**a**], purple: V7G [**b**], green: Y11A [**c**], orange: D9G [**d**], black: R15A [**e**], and blue: Y11P [**f**]) are overlapped with the same spectrum of 0.1 mM IscU WT (red). The black box indicates the K128 residues.

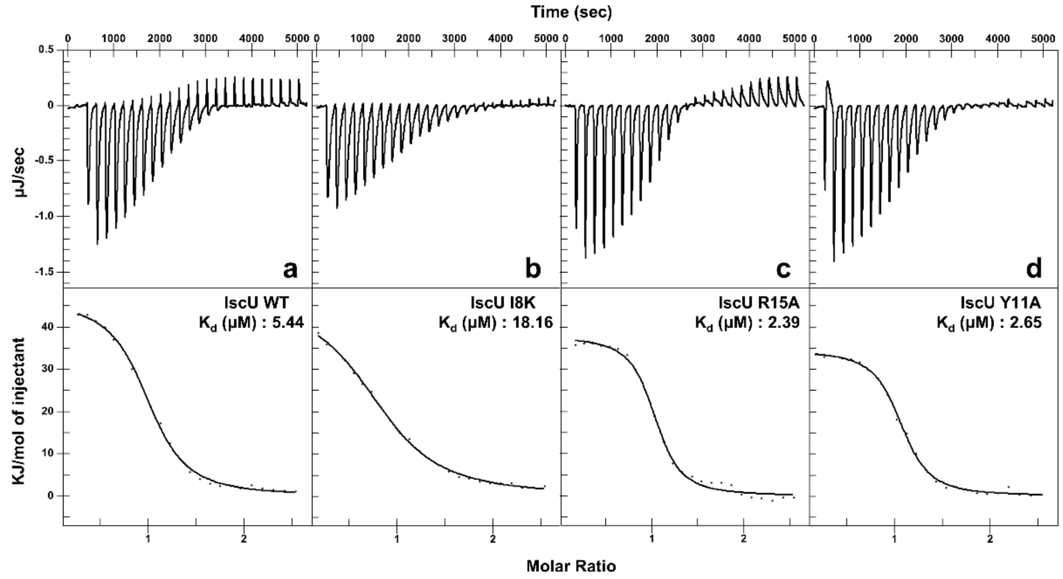


**Supplementary Fig 5. Isothermal titration calorimetry (ITC) results for the interaction of IscU mutants with HscA**

ITC results for the interaction of IscU WT (**a**) and the mutants, I8K (**b**), R15A (**c**), and Y11A (**d**), with HscA are shown with the calculated dissociation constants.

**Supplementary Table 1.** The SEC elution volume, and the CD-based α-helix composition and transition temperature calculation results for IscU Y3 mutants

| | Elution volume (ml) | α-helix composition (%) | Transition Temp. (℃) |
|---|---|---|---|
| WT | 12.40 | 22.0 | 37.88 ± 0.20 |
| Y3W | 12.86 | 22.3 | 38.69 ± 0.26 |
| Y3M | 12.62 | 22.3 | 38.57 ± 0.24 |
| Y3A | 12.35 | 20.6 | 36.00 ± 0.39 |

**Supplementary Table 2**. The SEC elution volume, and the CD-based α-helix composition and transition temperature calculation results for IscU α1-helix mutants

| | Elution volume (ml) | α-helix composition (%) | Transition Temp. (℃) |
|---|---|---|---|
| WT | 12.40 | 22.0 | 37.88 ± 0.20 |
| I8K | 12.87 | 22.3 | 41.08 ± 0.15 |
| R15A | 11.58 | 18.8 | 33.46 ± 0.02 |
| D9G | 12.14 | 19.7 | - |
| V7G | 12.05 | 19.4 | - |
| Y11A | 11.93 | 20.7 | - |
| Y11P | 10.81 | 12.5 | - |
